# Cross Sectional and Longitudinal Imaging Reveals Spatiotemporal Divergence in Morphogenesis and Cell Lineage Specification between *in-vivo* and *in-vitro* Mouse Embryo during Pre- and Peri-implantation

**DOI:** 10.64898/2025.12.15.691388

**Authors:** Huanhuan Yang

## Abstract

*In-vitro* culture is an essential step in assisted reproductive technology. Previous studies using sectional approaches have reported the impacts of *in-vitro* technologies on mammalian embryogenesis and offspring development. However, systematic and longitudinal comparisons between *in-vivo* and *in-vitro* spatiotemporal morphogenesis and cell lineage specification remain incomplete. Moreover, the phototoxicity of laser confocal microscopes on embryos is not fully evaluated. This study aims to bridge this gap using 4D live imaging via a laser scanning confocal microscopy to capture dynamic embryo development from a *CAG::H2B-GFP* mouse line. The results showed that *in-vitro* culture under a confocal microscopy cumulatively slowed the increase in embryonic cell number and embryos size over time. Furthermore, *in-vitro* embryo compaction, cavitation, primitive cell migration and hatching deferred in both spatial pattern and temporal progression. Additionally, *in-vitro* and *in-vivo* cell lineages differed in spatial distribution and expansion rate. These findings demonstrate that *in-vitro* culture delays embryo development, alters morphological events and disrupts cell lineage specification at both spatial and temporal scales during pre- and early peri-implantation. This work provides insights on the detailed extent of similarities and differences between *in-vivo* and *in-vitro* embryogenesis, beneficial when translating *in-vitro* findings into clinical, industrial and ecological applications.

## INTRODUCTION

Mammalian embryo development involves dynamic spatiotemporal coordination of multiple cell types at various levels, ranging from molecular, cellular aspects to cell population (Kojima *et al*., 2014; Niwayama *et al*., 2019; Fischer *et al*., 2020). *In-vitro* culture, essential for assisted reproductive technology (ART), and laser scanning confocal microscopes (LSCM) are widely used technologies to visualise and examine intricate spatiotemporal embryogenesis unfeasible *in vivo* (Yang, 2025).

To translate *in-vitro* findings to clinical practice, agriculture management and diversity conservation, it is essential to assess how closely experimentally cultured embryos, at all scales of possible analysis, replicate those developing naturally in the mammalian oviduct and uterus. Previous comparative studies between *in-vivo* and *in-vitro* preimplantation embryos, mainly from mouse, showed that *in-vitro* culture compromised cell cleavage, blastocyst formation, total cell numbers in blastocyst (Kim *et al*., 2004), hatching rate (Mahdavinezhad *et al*., 2019), and zona pellucida (ZP) properties (Coy *et al*., 2010). *In-vitro* cultured embryos also showed different patterns of gene expression, epigenetic levels, and transcriptomic/proteomic profiles (Doherty *et al*., 2000; Rinaudo and Schultz, 2004; Gardner and Lane, 2005; Gad *et al*., 2012; Bertoldo *et al*., 2015). However, these studies mainly employed sectional static approaches to analyse embryogenesis at single time points or used *in-vitro* fertilisation (IVF) or *in-vitro* oocyte maturation (IVM); static approaches may miss important developmental events and IVF/IVM procedures themselves can lead to changes in ZP, first cell cleavage, and molecule levels (Kim *et al*., 2004; Coy *et al*., 2010; Mahdavinezhad *et al*., 2019). In recent years, systematic updates of influences of *in-vitro* culture procedures on embryogenesis is lacking. Thus, it is essential to longitudinally visualise, track and investigate spatiotemporal morphological events and cell lineage specification in *in-vivo* and *in-vitro* embryos while minimising the influence of laboratory intervention and the effects of *in-vitro* culture on gametes when applicable. Furthermore, given the extensive use of LSCM in live embryo/cell imaging, it is also fundamental to examine the impact of LSCM visualisation-related technologies on embryogenesis.

In this study, aiming to enhance our understanding of the extent to which embryos under *in-vitro* culture conditions with LSCM live imaging reflect the natural dynamics of embryogenesis, I revisited and compared spatiotemporal features of mouse embryo development throughout pre-and early peri-implantation both *in vivo* and *in vitro*, using embryos from *CAG::H2B-GFPxCAG::H2B-GFP* (H2B-GFP) mouse and live-embryo imaging. This comparative study demonstrates *in-vitro* culture with live imaging impairs embryo growth, alters morphological events and disrupts cell lineage specification at both spatial and temporal scales during pre- and early peri-implantation.

## RESULTS

### Integrating Cross-Sectional and Successful Longitudinal Live Imaging into Early Embryo Development Analysis

To elucidate the intricate spatiotemporal dynamics of mouse embryo development *in vivo* and *in vitro*, I produced both cross-sectional and longitudinal data of H2B-GFP embryos. The cross-sectional data were collected from freshly flushed embryos at embryonic day (E) E1.0, E1.5, E2.5, E3.0, E3.5, E 3.75, E 4.5 and E4.75, serving as the *in-vivo* group (Fig 1Aa-c). The longitudinal data consisted of 4D videos capturing the developmental progression of embryos from 2-cell (E1.5) to the over 100-cell stage (∼E4.5/E4.75) through Nikon A1, serving as the *in-vitro* group (Fig 1Aa,Ab,Ad). Throughout this study, the terms “*in vivo*” and “freshly flushed embryos” were used interchangeably. The terms “*in vitro*” and “confocal microscope cultured embryos” were also used interchangeably. Both *in-vivo* and *in-vitro* embryos were sourced from at least three mouse litters (Table 1), except for midnight collections, which originated from a single litter due to practical constraints in the animal facility at The University of Manchester (UoM).

**Fig 1.**
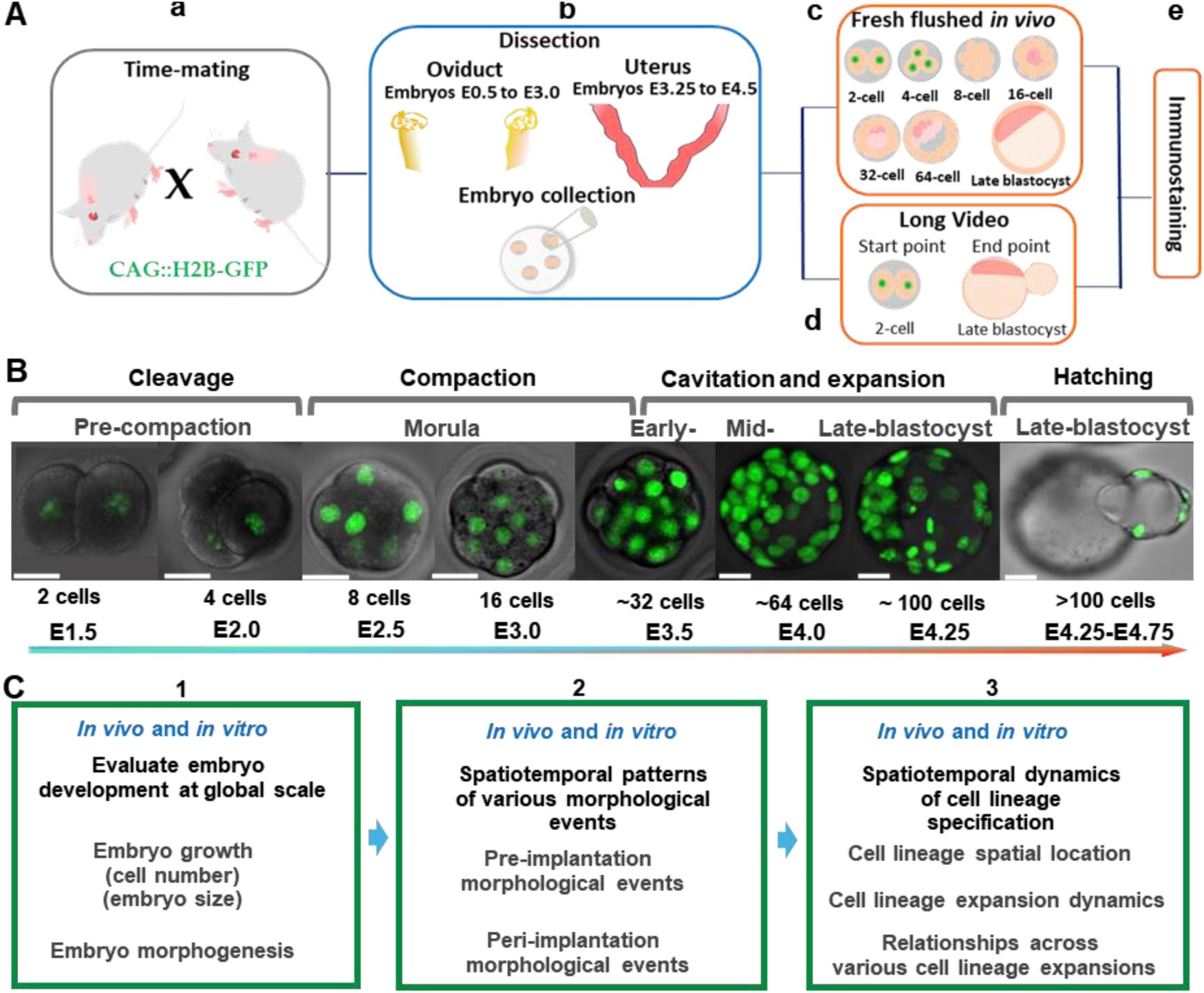
Workflow, data categorisation, and results structure. **(A)** Major experimental steps for collecting sectional and longitudinal data. **(B)** Embryo development and morphological events in long live videos imaged using a laser scanning confocal microscope (from E1.5 to E4.5, ∼66 hours), with developmental stages indicated with cell number and embryonic days. Scale bar: 20 μm. **(C)** Measured key parameters and the results layout—(C1) Cell number increase, embryo shape and size between the *in-vivo* and *in-vitro* groups; (C2) Spatiotemporal comparisons of morphological events during pre- and early peri-implantation stages between the two groups where feasible; (C3) Temporal and spatial patterns of *in-vivo* and *in-vitro* cell lineage expansion.

**Table 1.**
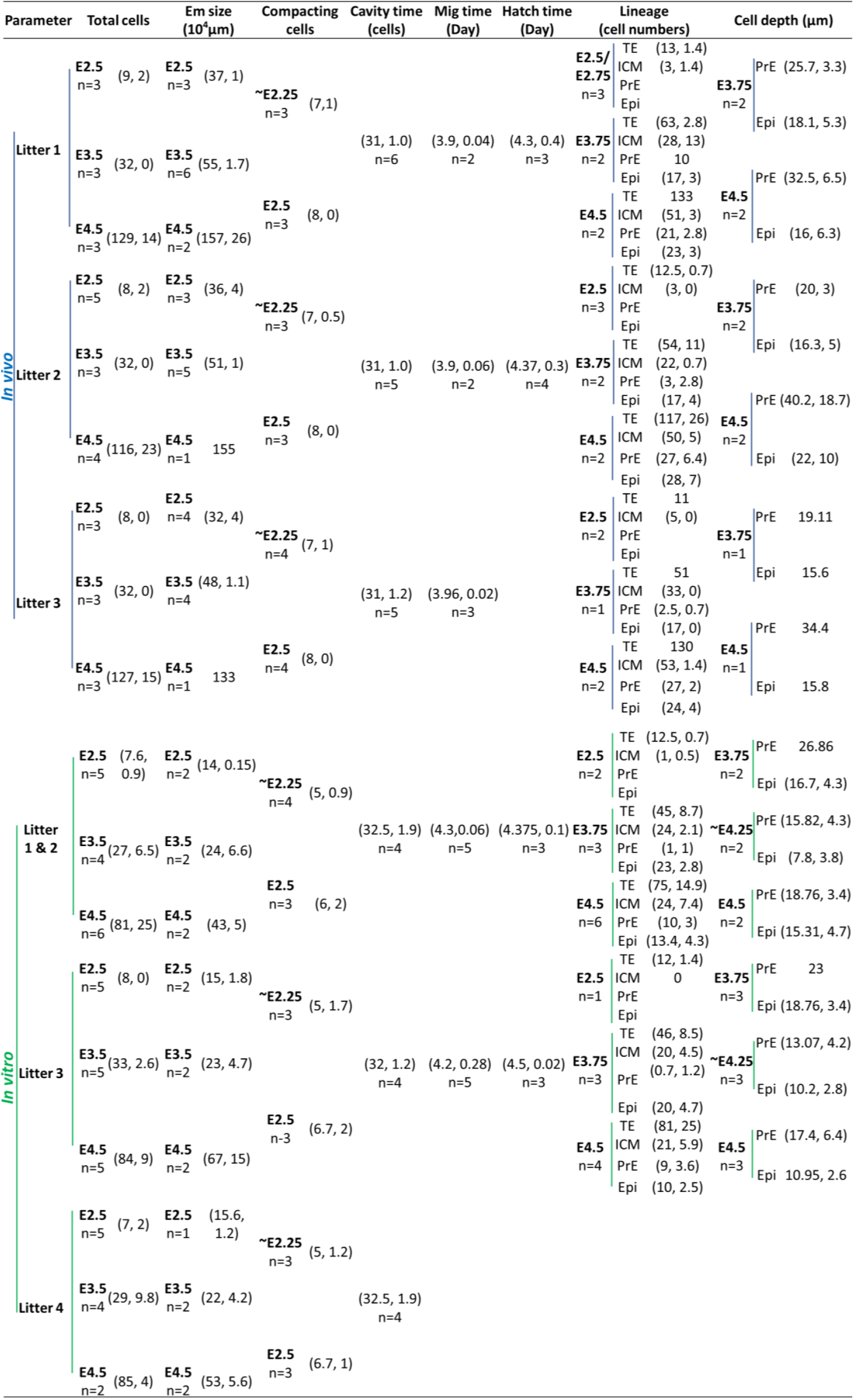
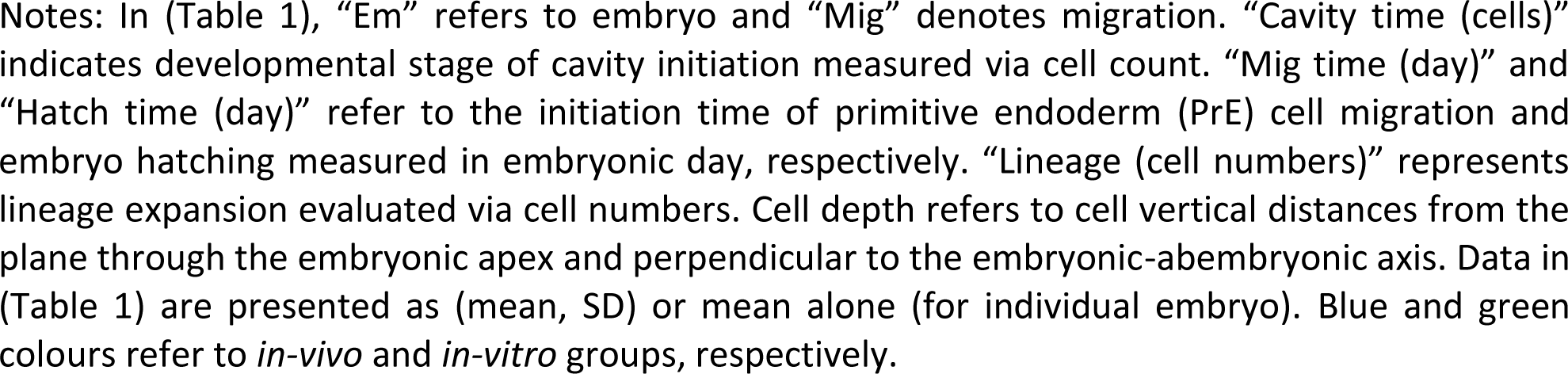
Variability of key parameters across embryos and litters *in vivo* and *in vitro*.

Non-invasive long time-lapse videos successfully exhibited pre- and early peri-implantation embryo development over E1.5 to E4.5/E4.75 (∼65 hours). In three 4D long videos with 36 embryos (from 3 litters), only one embryo, whose development was arrested at the 2-cell stage, was excluded from subsequent analyses. The remaining embryos displayed typical developmental trends, including cleavage, compaction, cavitation and hatching (Fig 1B). Attempts were also made to record long videos from zygotes using LSCM. However, the majority (16 out of 23) of these zygotes experienced mortality or abnormal development and thus were not included here.

These experiments demonstrate that a 2-cell embryo can develop continuously to the ∼100-cell stage *in vitro* under LSCM, allowing for a comparison of *in-vitro* embryo growth, morphogenesis and cell lineage expansion with *in-vivo* counterparts (Fig 1A-C).

### Initial Evaluation of Key Parameters across Mouse Litters and Embryos

Key aspects of embryo morphogenesis and lineage specification were first examined across mouse litter and among embryos within each litter, both *in vivo* and *in vitro*. These parameters included embryonic cell number growth at each stage, changes in embryo size in 3D over time, timing of embryo compaction and initiation of migration and hatching, and cell lineage expansion (Table 1). Morphological event timing was assessed by cell counts. Descriptive statistics are presented as (mean, SD). A Kruskal-Wallis test was performed to assess variability across litters (where n ≥ 3 embryos/litter), and the results indicated no statistically significant differences across litters in the evaluated parameters in either *in-vivo* or *in-vitro* group. This initiation analysis confirms that embryos from different litters of the H2B-GFP mouse line generally share a consistent baseline, validating the homogenous foundation for further analyses detailed in the following sections.

### Temporal and Spatial Dynamics of Embryo Growth Both *In Vivo* and *In Vitro*

#### Temporal trends of cell number increase

Cells in both *in-vivo* and *in-vitro* embryos were counted to assess temporal changes in embryo growth (see Materials and Methods for the cell-counting approach), with sample sizes shown in (Fig 2A). The results showed that cell numbers increased more exponentially over time in the *in-vivo* group compared to the *in-vitro* one, which exhibited a slower and more irregular rate of cell number increase (Fig 2B). Both the *in-vivo* and *in-vitro* groups exhibited a slightly faster increase in cell numbers after the 8-cell stage, with a sharp rise around E3.5 (Fig 2B). The gradually increased differences in cell number between the two groups were statistically significant at E3.75, E4.5, and E4.75 (Mann Whitney test, *p* = 0.031, *p* < 0.0001, and *p* = 0.036, respectively) (Fig 2B). Taking cell number as a reference to examine development timing, *in-vitro* embryos exhibited remarkable developmental delays compared to the *in-vivo* embryos, with delays of 1.5 to three hours after E3.25 and over six hours around E3.75, extending up to 12 hours until the peri-implantation stage (Fig 2B).

**Fig 2.**
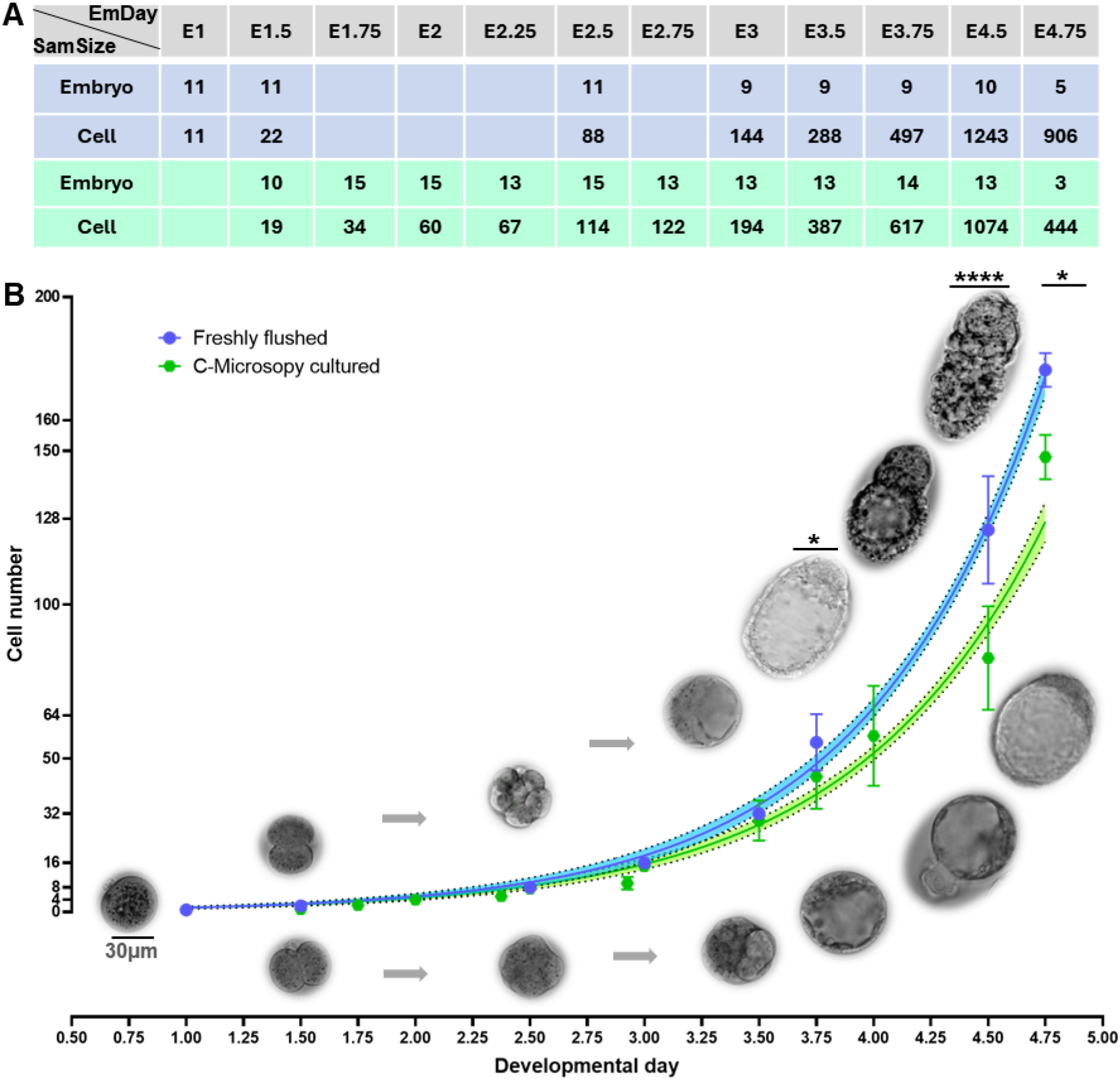
Embryonic cell number changes over developmental days. **(A)** Sample sizes of embryos and cells analysed in *in-vivo* (blue) and *in-vitro* (green) embryos, with “EmDay” representing embryonic day and “SamSize” for sample size. **(B)** Embryonic cell numbers *in vivo* (blue) and *in vitro* (green) over developmental stages, with non-linear curve fit results of *in-vivo* (“Freshly flushed”) and *in-vitro* (“CM cultured”-embryos cultured under the confocal microscope) cell counts, along with 95% CI (profile likelihood) (dashed areas) and SD error bar. *In-vivo* and *in-vitro* embryo morphologies are presented at the top and bottom of the blue and green curves, respectively, with grey arrows for embryos having the same cell numbers on the same developmental days but with different morphogenesis. Scale bar: 30 μm. Mann Whitney test was applied for cell numbers comparison in both groups on each developmental day, with \**p* < 0.05, ** *p* < 0.01, \*\*\*\**p* < 0.0001. *In-vivo* embryos were from three litters, and *in-vitro* ones were from three videos across four litters.

These data show that *in-vitro* cultured embryos under LSCM exhibit reduced embryo growth rates, with delays in cell number increase becoming significant from E3.75, highlighting the influence of *in-vitro* conditions on early embryonic development.

### Spatiotemporal trends of embryo shape and size *in vivo* and *in vitro*

Embryo shapes differed between the *in-vivo* and *in-vitro* groups, despite similar cell numbers on certain developmental days (Fig 2B, grey embryos). Disparities in embryo shape between *in-vivo* and *in-vitro* groups started to show around E2.5. *In-vivo* embryos appeared to initiate compaction slightly earlier than their *in-vitro* counterparts. From E2.5 and E4.5, *in-vitro* morphological development continued slowing down compared to *in-vivo* group (Fig 3A,B). Before compaction, embryos in both groups appeared as loose clusters, with cells displaying predominantly round or slightly ellipsoid and distinguishable cellular boundaries (Fig 3A,B). Post-compaction, cells showed diverse shapes in both groups, adhering tightly to each other, in line with previous studies (Ducibella *et al*., 1975; Niwayama *et al*., 2019), resulting in a cohesive and integrated developmental architecture (Fig 3A,B). Around E3.5-E4.0, the 12 observed *in-vivo* embryos became ellipsoid, contrasting with the round shape retained by 94% of 34 *in-vitro* embryos (Fig 3A,B). Two of the 34 *in-vitro* embryos exhibited a slight ellipsoid shape post-ZP breakage. During early peri-implantation (E4.25-E4.5), *in-vivo* embryos elongated along the embryo-abembryonic axis, while *in-vitro* embryos sustained their round or slight ellipsoid configuration (Fig 3A,B).

**Fig 3.**
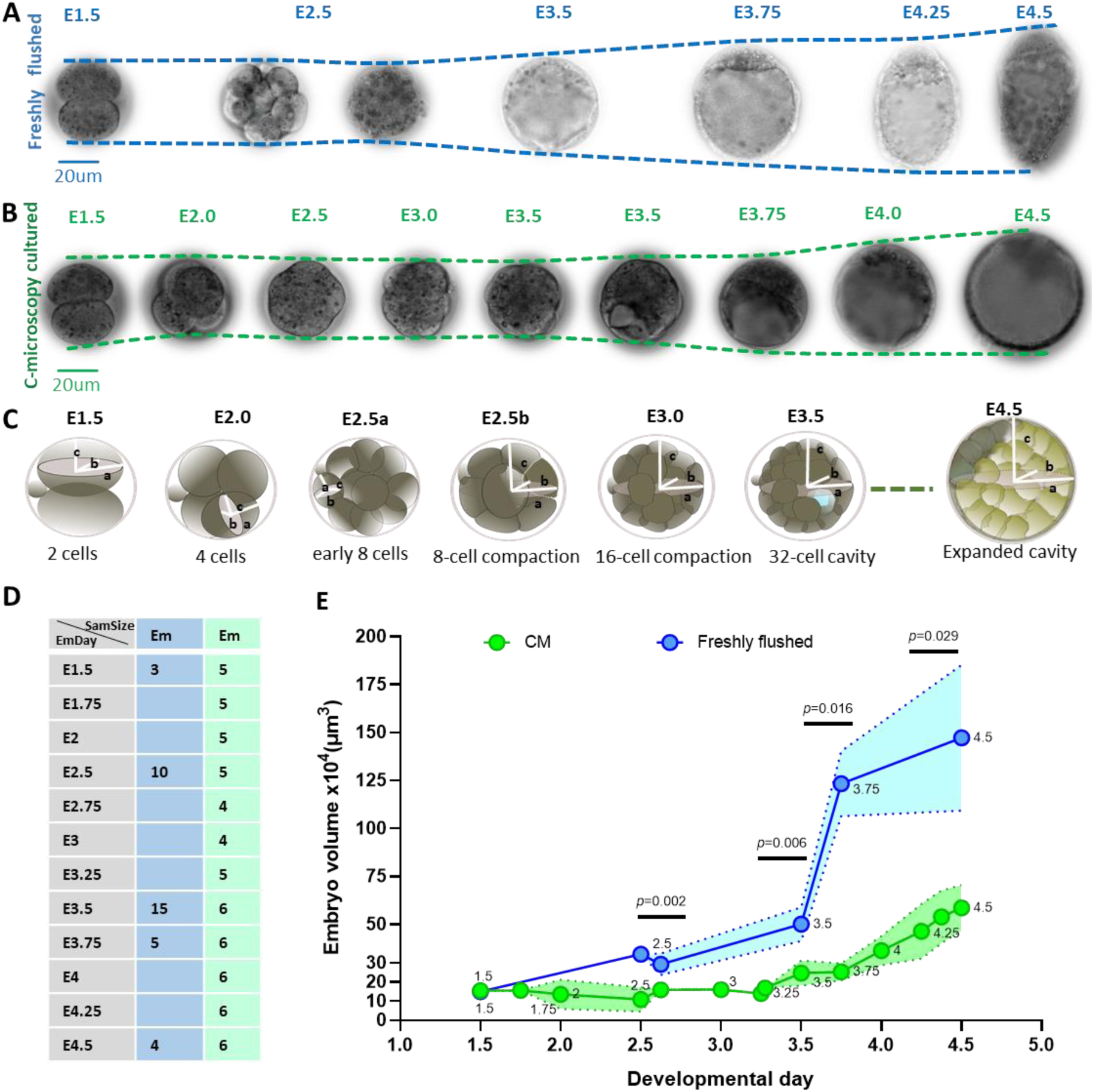
Embryo morphology and size changes *in vivo* and *in vitro* during pre- and per-implantation stages. **(A,B)** Embryo growth and morphological changes *in vivo* (blue) and *in vitro* (green), with dashed lines linking apexes of embryonic (top) and abembryonic (bottom) parts at each developmental stage. Scale bar: 20 μm, and variation in grey shades reflecting different presentation brightness. **(C)** Methods for measuring embryo size, with a, b and c for semi-major, semi-intermediate and semi-minor axes of ellipsoid cells before compaction and ellipsoid embryos post-compaction, as detailed in Materials and Methods. **(D)** Sample sizes *in vivo* (blue) and *in vitro* (green), with “Em” indicating embryo, “EmDay” referring to embryonic day and “SamSize” representing sample size. **(E)** Man-Whitney test results of embryo size changes from E2.5 to E4.5 *in vivo* (blue) and *in vitro* (green), with “Fresh flushed” for the *in-vivo* group and “CM” for the *in-vitro* group, shaded areas between dashed lines for 95% CI of original data. *In-vivo* embryos were from three litters, and *in-vitro* ones were from two videos across two litters.

Informed by observed geometric attributes of cellular and embryonic shapes, embryo size was measured over time in both *in-vivo* and *in-vitro* groups (Fig 3C-E; see Materials and Methods for details on embryo size measurement). The sample sizes are shown in (Fig 3D). The embryo size assessment curve showed faster growth of *in-vivo* embryos than *in-vitro* ones, particularly after E3.5 (Fig 3E). Embryo volume in both groups was similar at E1.5. *In-vivo* embryo volume sharply increased between E3.5 and E3.75, and then slowed down between E3.75 and E4.5 (Fig 3E). *In-vitro* embryo volume increased rapidly between E3.5 and E4.25, and slowed down by E4.5. Embryo size changes showed statistically significant differences between *in-vivo* and *in-vitro* groups from E2.5 (Mann-Whitney test, see Fig 3E for detailed *p* values). Despite increased cell numbers, *in-vitro* embryo size displayed slight drops at E2.5 and E3.25, coinciding with embryo compactions (Fig 3E). Furthermore, a minor decrease was observed at E3.0, mirroring the timing of 16-cell morula formation (Fig 3E). *In vivo*, due to limited access to embryo collections between E1.5 and E2.5, size measurements were taken only around E2.5 and E3.5, leading to a single detectable size decline after E2.5 (Fig 3E).

These data demonstrate that *in-vitro* embryos exhibit significantly slower growth in embryo size from E2.5 onwards, further emphasising the impact of *in-vitro* conditions on embryo development.

### Spatiotemporal Patterns of Morphological Events *In Vitro* and *In Vivo*

Based on the above-mentioned findings, I investigated the spatial patterns and temporal trends of consecutive key morphological events, including compaction and cavitation during pre-implantation, as well as migration and hatching during early peri-implantation, within both *in-vitro* and *in-vivo* groups (see Materials and Methods for detailed measurements).

### Spatiotemporal features of compaction and cavitation

For pre-implantation embryos, I first investigated spatial and temporal patterns of compaction, a process crucial for cell lineage specification. *In-vitro* video in this study showed that compaction occurred gradually from initiation to completion. Thus, different phases of compaction were classified to further analyse this process. The initial attachment was defined as the first observed site of tight cell-cell contact, characterised by a cell morphology transition from a spherical to a flattened shape within a localised cell region, leading to uneven contours along the cell borders (Fig 4Aa,Ab). Multiside attachment was evaluated as the presence of more than one cell-cell contact site along the edges of flattened cells (Fig 4Aa,Ab). Complete compaction was reached when all cells were flattened and tightly adhered to each other, with cell borders becoming indistinguishable (Fig 4Ac).

**Fig 4.**
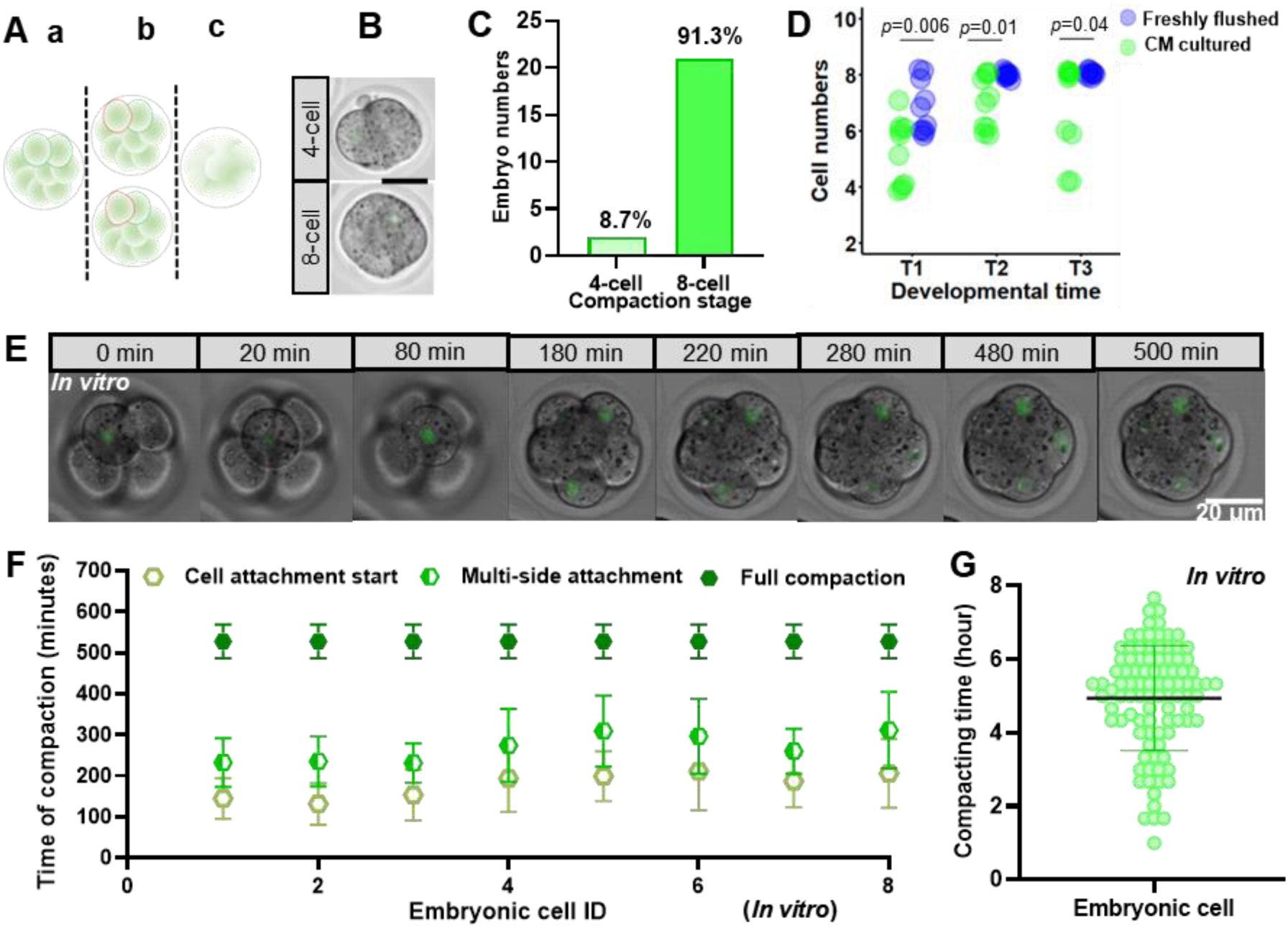
Spatial patterns and timing trends of 8-cell embryo compaction *in vivo* and *in vitro*. **(A)** Local cell attachment detection *in vitro*, with geometrical shape of blastomeres without visible attachment (green circled cells) in (Aa), initial attachment–the first site of tight cell-cell contact between neighbouring cells with shape changes from spherical to flattened (red circled cells in the top embryo), multiside attachment–multiple cell-cell contact sites/areas along the edges of the flattened cells (red circled cells in the bottom embryo) in (Ab), and complete compaction with all blastomeres showing indistinguishable cell borders due to tight cell-cell contact in (Ac). **(B)** Spatial compaction patterns *in vitro*. **(C)** The occurrence of different compaction patterns *in vitro*. **(D)** Attachment timing differences *in vivo* (blue) and *in vitro* (green) and Man-Whitney test results at T1 (embryos collected between 8:00 and 9:00 am), T3 (collected at E4.5-noon), and T2 (embryos collected between T1 and T3). Sample size: 10 embryos (three litters) at each stage. **(E)** Morphology changes during compaction from initial attachment to complete compaction. **(F)** The timing of various attachment phases of eight cells per embryo during compaction progression *in vitro* (10 embryos from 3 litters). Yellow stands for initial cell attachment, with green for multiple-side attachment, and dark green for complete compaction. In green dark plot, identical timing and error bars show that all cells within each embryo simultaneously reached full compaction phase, reflecting the definition of complete embryo compaction in this study (all blastomeres contacted each other tightly and flattened with indistinguishable cell borders). X-axis represents embryonic cell IDs (defined by their division order) per embryo. **(G)** Duration of *in-vitro* compaction from initial attachment to completion for each 8-cell blastomere across 12 embryos from three litters, with SD error bar. Scale bar in (B,E): 20μm.

The results showed that 8.7% of *in-vitro* embryos (n=23) initiated compaction at the 4-cell stage and 91.3% at the 8-cell stage, both occurring around E2.5 and progressing to the late blastocyst stage (Fig 4B,C). However, embryos compacting at the 4-cell stage experienced delays in subsequent developmental events such as cavitation. *In-vivo* embryos (n=23) had already shown local compaction before E2.5 when they were collected, with more cells attached at least 4 hours significantly earlier than the *in-vitro* group (Fig 4D). *In-vivo* embryos compacted at the 4-cell stage were rarely observed within embryo collection timeframes. It is noteworthy that exploring embryo compaction dynamics *in vivo* is challenging due to the difficulty in continuously observing *in-vivo* compaction without invasive techniques and disrupting the natural environment of embryo development. Therefore, I opted to examine the *in-vitro* group, focusing on the temporal features of the predominant compaction pattern at the 8-cell stage. The results demonstrated that embryonic cells were locally attached to neighbouring cells asynchronously. Specifically, attachments occurred at different times both between cells and across different sides of individual cells, over 4.36 ± 1.53 hours until complete compaction (Fig 4E-G).

Following compaction, I examined the spatiotemporal patterns of embryo cavitation *in vivo* and *in vitro*. Cavitation initiation was identified by the first detection of a small clear or translucent area (micro lumens) within the embryo, observed under a confocal and light microscopes. *In vitro*, among 19 embryos, three cavitation patterns were detected: 57.89% showed a single cavity, 21.05% showed multiple merged cavities at similar regions, and 21.04% showed independent multiple cavities at various sites (Fig 5Aa-c,Ba). In the latter multiple cavities, only one major cavity remained and expanded while the others collapsed over time (Fig 5Ab). These multiple cavities rarely led to the separation of inner cell mass (ICM) into two or more compartments, occurring in only one of the latter embryos. *In vivo*, across 21 embryos collected around E3.25, 52.4% already had displayed a single cavity pattern, 14.3% exhibited multiple cavities in similar regions within embryos, and 33.3% had not initiated cavitation as early as E3.25 (Fig 5Bb). Around E3.5, most *in-vivo* embryos exhibited expanded cavities, with only 7% of 54 embryos displaying small cavities, probably indicative of cavity initiation or delayed development. *In-vitro* cavitation initiation spanned from ∼E3.25 to ∼E3.5, with 50%, 33.3% and 16.7% of 12 embryos initiating at the 31-cell, 32-cell or 35-cell stage, respectively (Fig 5C, green). *In-vivo* cavitation initiation among 14 embryos around E3.25, where embryos showed small cavities, occurred before the 30-cell (35.7%), 31-cell (42.9%) and 32-cell (21.4%) stages (Fig 5C, blue). The cell number-staged timings of cavity initiation significantly differed between both groups (Mann-Whitney test, *p* = 0.0217).

**Fig 5.**
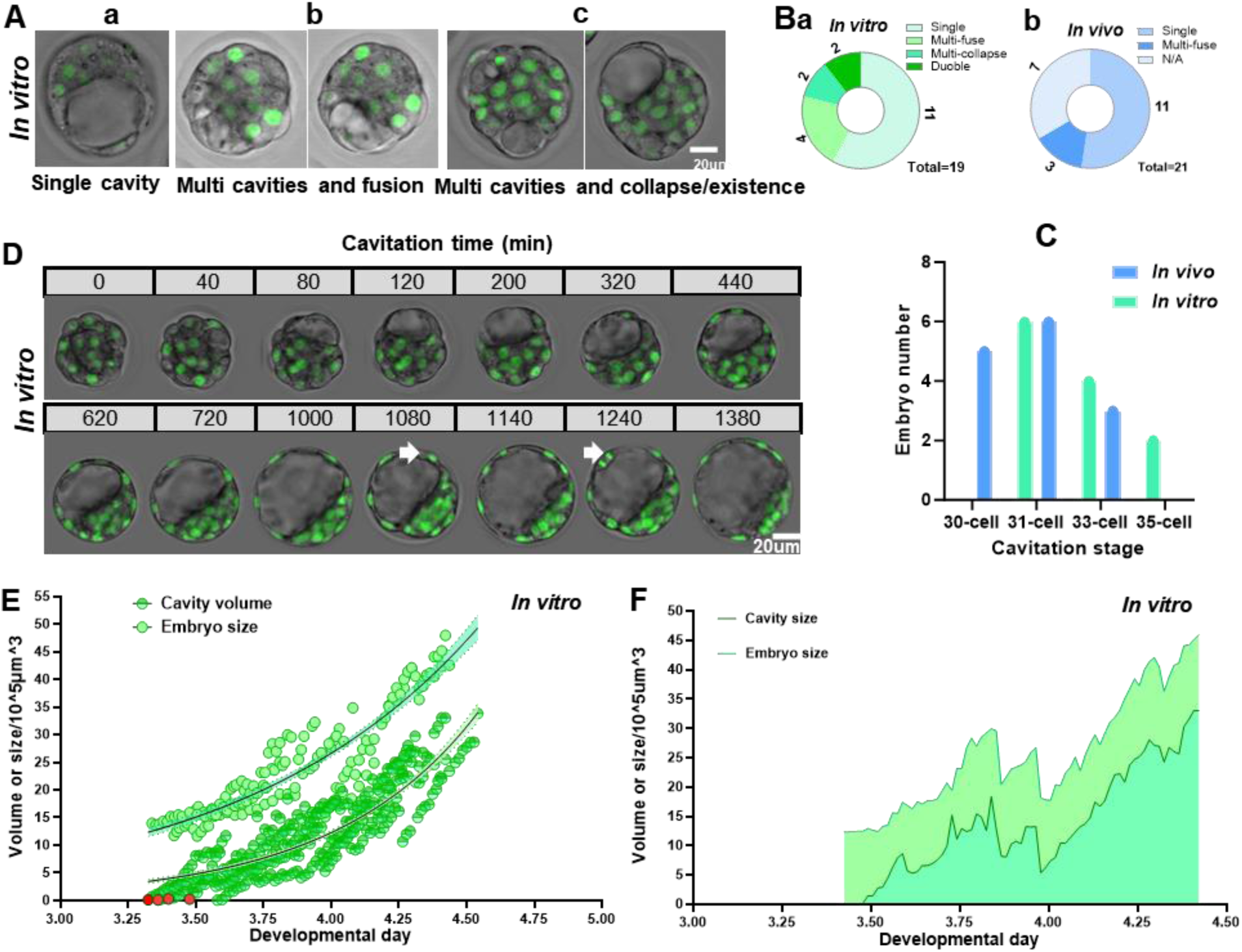
Spatial patterns, time trends, and expansion features of embryo cavitation. **(A)** Various patterns of embryo cavitation initiation. **(B)** Distribution frequency of cavitation initiation patterns *in vivo* (green) and *in vitro* (green). The sample size of analysed embryos: 21 (three litters) for the *in-vivo* group and 19 (four litters) for the *in vitro* group. In (A,B), “Multi” for “multiple”. **(C)** Timing of cavitation occurrence *in vivo* (14 embryos) and *in vitro* (12 embryos), using cell number as a reference. **(D)** Cavity morphological changes over time, including cavity collapse. White arrows point at dividing mural TE cells. Scale bar in (A,D): 20μm. **(E)** Non-linear curve fitting results of embryo size changes and cavity expansion over time, with red dots indicating the timing of cavitation initiation referencing developmental day. Sample size: five embryos from two litters. **(F)** Cavity expansion and embryo enlargement over time within individual embryos, with areas between curve lines and the X-axis representing cumulative embryo size (light green) and cavity volumes (dark green).

The continuous changes in cavities *in vitro* were investigated, showing progressive expansion in all analysed embryos (Fig 5D). However, around the late blastocyst stage (E4.0-E4.5), 16 out of 19 embryos experienced considerable reductions in cavity volume, followed by re-expansion, typically around 1080 and 1240 minutes post-cavity initiation (Fig 5D). Four embryos underwent cavity reduction at earlier stages, around E3.75, coinciding with certain mural trophectoderm (TE) cell divisions (Fig 5D, white arrows). The quantitative analysis of approximate cavity volume relative to embryo size over time showed that cavity expansion generally exhibited exponential growth patterns (Exponential growth model, R^2^ = 0.96), particularly during the late blastocyst stages, accounting for over 60%-70% of embryo size (Fig 5C,E,F; see Materials and Methods for cavity volume measurement). At the individual embryo level, cavity expansion demonstrated irregular growth speed (Fig 5F). The speed of cavity expansion decreased around E4.25 to E4.5, resulting in slow increase in cavity volume and embryo size (Fig 5F). After E4.5, *in-vitro* cultured embryos consistently encountered collapse, leading to minimal cavity area and failure to re-expand in glass bottom dishes, causing notable morphological differences from *in-vivo* embryos at the same stage.

Overall, the present findings demonstrate that embryo compaction is a gradual process, and *in-vitro* embryos show slower compaction than *in-vivo* ones. Furthermore, my findings suggest that *in-vitro* embryos initiate cavitation later, at higher cell numbers, and experience notable collapse around E4.5. These results suggest the effects of *in-vitro* conditions on both the spatial pattern and temporal progression of the morphological events of pre-implantation embryos.

### Spatiotemporal features of PrE cell migration and embryo hatching

During the peri-implantation stages, the earliest event observed both *in vivo* and *in vitro* was cell migration, identified using multiple IMARIS viewer tools to track the sustained movement of cell nuclei, marked by H2B-GFP, from E3.5 to E4.5. One to three cells per observed embryo showed sustained displacement from the deep ICM or central ICM surface to the peripheral ICM surface. Their nuclei positions extended beyond the contiguous boundaries of adjacent ICM surface cells, confirming directed movement. These cells exhibited protrusions (Fig 6A), as previously described (Enders *et al*., 1978). The observation of cell protrusions supplemented nuclei position tracking to support cell migration assessment (Fig 6A). To delineate this phenomenon, I introduced the term “migrating pioneer cells” to characterise the initial subset of one to three cells with protrusions that initiated the migration process (Fig 6Aa,Ab). These *in-vivo* and *in-vitro* migrating pioneer cells appeared at varied times across various peripheral areas along the ICM surface layer (PrE cell population), indicating an asynchronous spatiotemporal pattern of PrE migration. Significant disparities were observed in both cell number- and embryonic day-specified timings of migration initiation between *in-vivo* and *in-vitro* groups (Mann Whitney test, *p* = 0.045 and *p* = 0.0008, respectively) (Fig 6B,C). *In vivo*, embryos had exhibited migration pioneer cells featuring protrusions around the 70-cell stage (70 ± 8 cells) by ∼E4.0, while *in vitro*, migrating pioneer cells were observed later, at around the 82-cell stage (83 ± 12 cells) at E4.25-E4.5 (Fig 6B,C). Note that, as protrusive activity alone does not always lead to full migration, distinguishing true cell movement from transient protrusion events remains a limitation of this approach to some extent.

**Fig 6.**
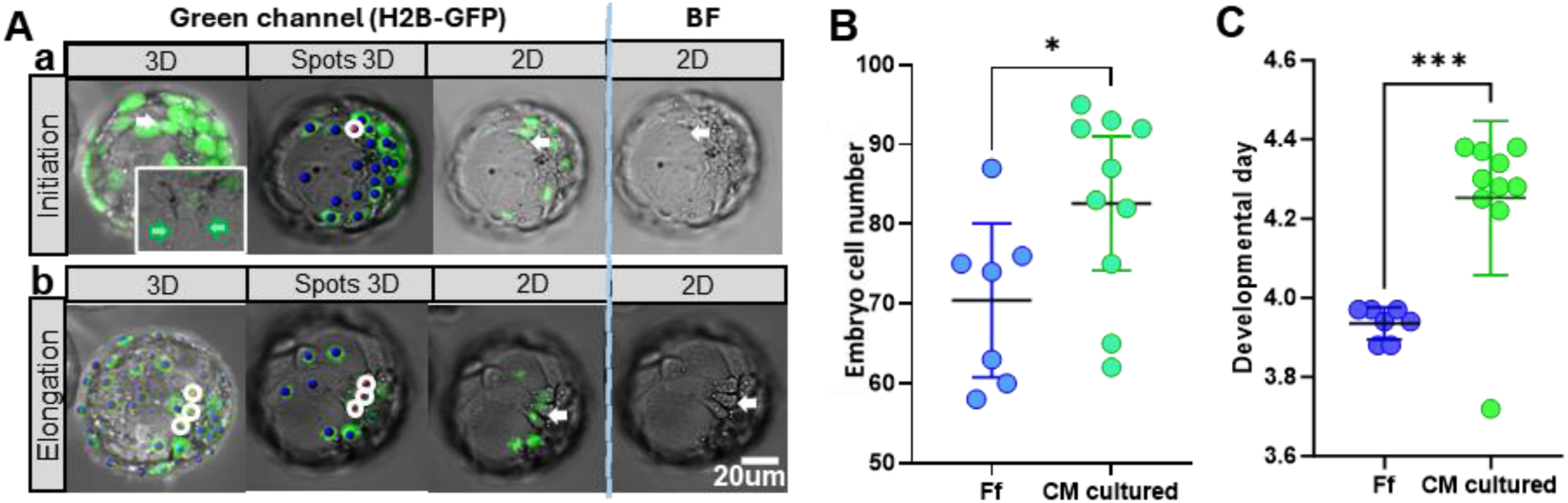
Morphologies of migrating pioneer cells and timing of migration event *in vivo* and *in vitro.* **(Aa,Ab)** Changes in protrusion morphology of migrating PrE cells over time via green channel and bright field (BF) in 3D and 2D views. White arrows indicate migrating pioneer cells, with protrusions denoted by green arrows in the enlarged 3D viewer. Blue and red spots highlighted by white circles, added via IMARIS, confirm cell positions. Scale bar: 20μm. **(B)** Total embryo cell counts at the observation of the first migration cells featuring protrusions. **(C)** The timing of migration initiation in embryonic developmental day. In both (B,C), Mann Whitney test was employed. Error bars: SD. “Ff” denotes freshly flushed, mirroring the *in-vivo* group, and “CM” reflects the *in-vitro* group. Statistical significance: *p* = 0.05, \**p* < 0.05 and \*\*\**p* < 0.001. The embryo sample size: Seven (three litters) for the *in-vivo* group and 10 (three videos across four litters) for the *in-vitro* group.

The second notable morphological transformation during the peri-implantation stage observed *in vitro* was embryo hatching, although this was not always the case. The first few TE cells breaking free from the ZP during hatching are termed here “hatching breaker cells”. These hatching breaker cells passed through a slight crack within the ZP, subsequently followed by a gradual hatching of other cells from the ZP (Fig 7A,B). Hatching breaker cells were observed in different TE areas, including both mural TE (Mu-TE) and polar TE (P-TE) regions. To classify these hatching patterns, I segmented the embryo structure into four distinct 3D anatomical areas, using the ICM cap as the reference point for positive longitudinal directionality and the major hatching site as the positive horizontal direction. These segmented anatomical regions were denoted as I, II, III, and IV quadrants (Fig 7Aa,Ab). In quadrant I, 15.8% of 19 examined embryos hatched from the P-TE, while 84.2% of embryos hatched from the Mu-TE near the P-TE in quadrant IV (Fig 7A,C). Noticeably, three embryos underwent multi-hatching patterns, initially hatching from quadrant IV and subsequently from quadrant I, likely due to embryo crowding in culture dishes, indicating the impact of limited growth space exerted by the adjacent embryos on embryo hatching patterns. Additionally, two embryos showed hatching initiation in the centres of abembryonic parts, but hatching was either retracted or kept the initiation status during the subsequent development. Temporal feature results showed that *in-vivo* hatching occurred approximately three to four hours before E4.5 (96-cell stage) and *in-vitro* around E4.5 (the 98-cell stage) (Fig 7D,E). I then examined the hatching duration and speed, from the timing of the emergence of hatching breaker cells to the time when embryos fully exited the ZP. The results showed that hatching from quadrant IV took one to 2.5 hours, while embryos hatching from quadrant I were majorly trapped within the ZP (Fig 7F,G). Despite the similar timings of initial hatching based on cell number and developmental day in both *in-vivo* and *in-vitro* groups, there was a striking contrast in the hatching speed and outcomes between the two groups. *In vivo*, 99% of the 45 embryos had completed hatching before E4.5. However, *in vitro*, only 35% of 36 embryos were either in the middle of hatching or completed hatching at E4.5 stages. The remaining embryos had not initiated hatching at E4.5 or were trapped in the ZP, as described earlier.

**Fig 7.**
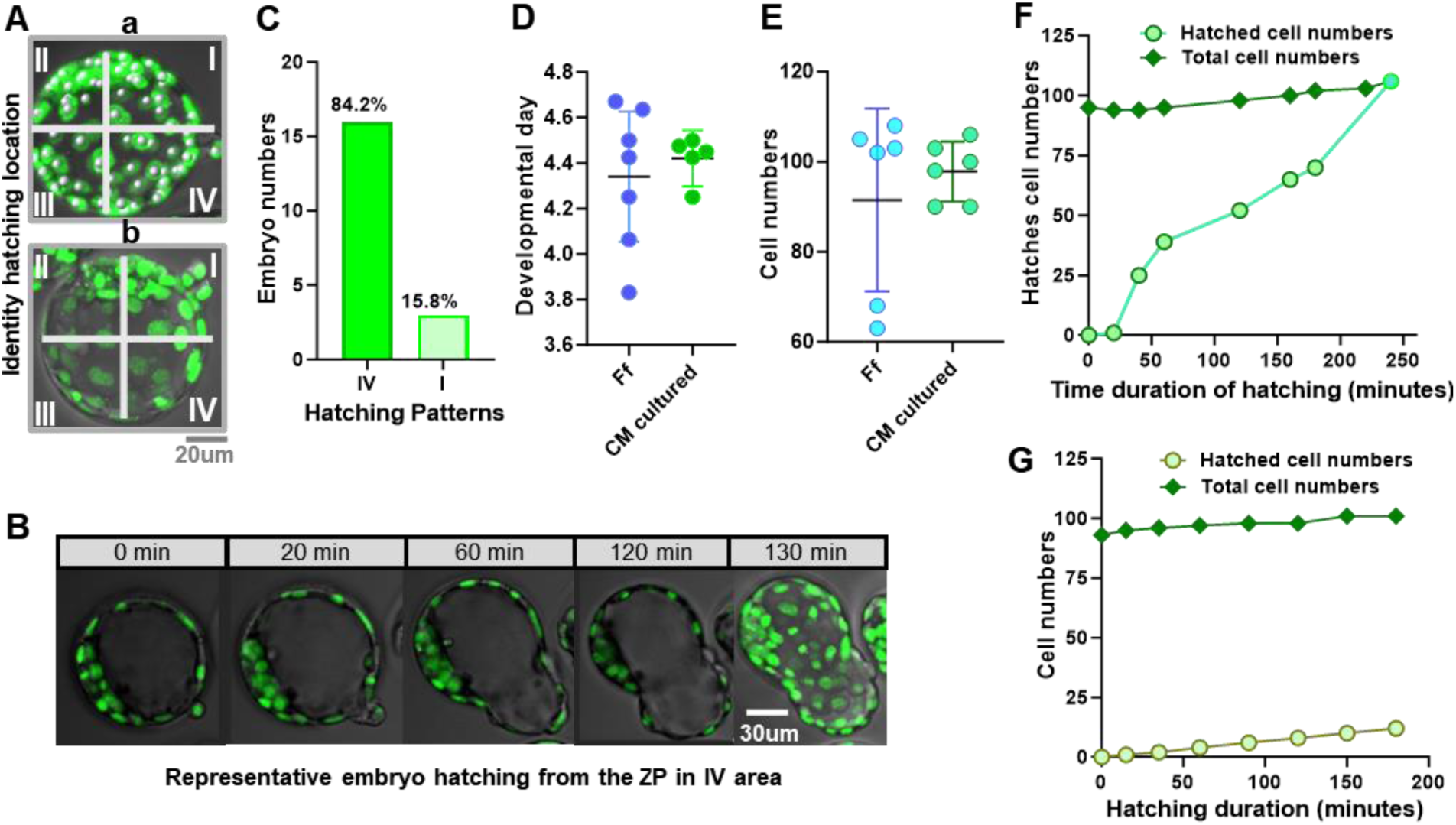
Spatial patterns and timing of embryo hatching *in vivo* and *in vitro*. **(A)** Different spatial sites of embryo hatching. Quadrants I, II, III, and IV denote anatomical areas based on hatching sites: quadrant I around P-TE, while IV was located in Mu-TE but not in the centre of abembryonic sites. Scale bar: 20μm. **(B)** Morphological changes during the hatching process. Scale bar: 30μm. **(C)** Distribution frequency of different spatial patterns of embryo hatching. **(D,E)** Hatching timing with SD error bar, referenced by developmental day and cell number, respectively. “Ff”, *i.e.*, freshly flushed, denotes the *in-vivo* group and “CM” stands for the *in-vitro* group. Sample size of analysed embryos: 19 (three litters) in (C). Seven for the *in-vivo* group and five-six for the *in-vitro* group (three litters) in (D,E). **(F,G)** Hatched cell numbers over time from quadrants IV and I in example embryos, respectively, with green for cells undergoing hatching and dark green for total embryonic cell numbers.

All in all, these findings illustrate that *in-vitro* embryos initiate migration later, at higher cell numbers, and exhibit slower and delayed hatching, especially at P-TE sites. These findings emphasise the impact of *in-vitro* conditions on the initiation and progression of peri-implantation morphological events.

### Spatiotemporal Trends of Cell Lineage Specification *In Vivo* and *In Vitro*

The notable spatiotemporal differences observed and highlighted above in embryo growth and morphological events *in vivo* and *in vitro* necessitated confirming whether there were corresponding spatiotemporal differences in protein expression of the cell lineages such as TE, PrE and epiblast (Epi) in both groups. To address this, *in-vitro* embryos from the E1.5 (2-cell stage) were cultured and recorded at different developmental stages from E2.5 to E4.5. Likewise, fresh embryos for the *in-vivo* group were collected and fixed accordingly. Immunofluorescence staining, using Sox2 and Gata4 markers, with Hoechst staining for cell nuclei, was employed to distinguish the Epi and PrE cell fates in both groups (refer to Materials and Methods for details of immunofluorescence staining). Both qualitative and quantitative analyses were conducted as detailed below.

### Qualitative observation of Sox2- and Gata4-cell spatiotemporal distribution

Immunostaining displayed similar temporal trends in Sox2- and Gata4-presence between *in-vivo* and *in-vitro* embryonic cells during E2.5 and E3.5 (Fig 8A,B). At E2.5 and E2.75 in both groups, Sox2 was detected in the nucleus, around the nuclear membrane, or in the cytoplasm of embryonic cells (Fig 8Aa-b,Ba-b). By E3.5, Sox2 became majorly restricted to the ICM cell nuclei, occasionally around the nuclear vicinity in certain ICM and TE cells (Fig 8Ac,Bc). At E3.5, E4.375 (*in vitro*) and E4.5, Sox2 was specifically limited to the ICM cell nuclei in both groups (Fig 8Ac-e,Bc-f). The staining of Gata4 encompassed the cell nuclear membrane at E2.5, E2.75 and E3.5 in both *in-vivo* and *in-vitro* settings (Fig 8Aa-c,Ba-c). At E2.75, Gata4 was visible in the nuclei of internal and external cells in certain *in-vivo* embryos (Fig 8Ab). At E3.5, Gata4 became less pronounced in most *in-vivo* embryonic cells, except for TE and a few ICM cells where it was still detected around the nuclear membrane (Fig 8Ac). *In-vitro*, Gata4 consistently encircled the nuclear membrane of all/most embryonic cells at E3.5 (Fig 8Bc). At E3.75, *in-vivo* embryos exhibited 3 to 4 Gata4-positive (in the nuclei) cells, while *in-vitro* embryos had 0 to 1 Gata4 cell, which began to appear noticeably around E4.375 (Fig 8Ad,Bd,Be). By E4.5, more expanded Gata4- and Sox2-positive cells were observed *in vivo* than *in vitro*.

**Fig 8.**
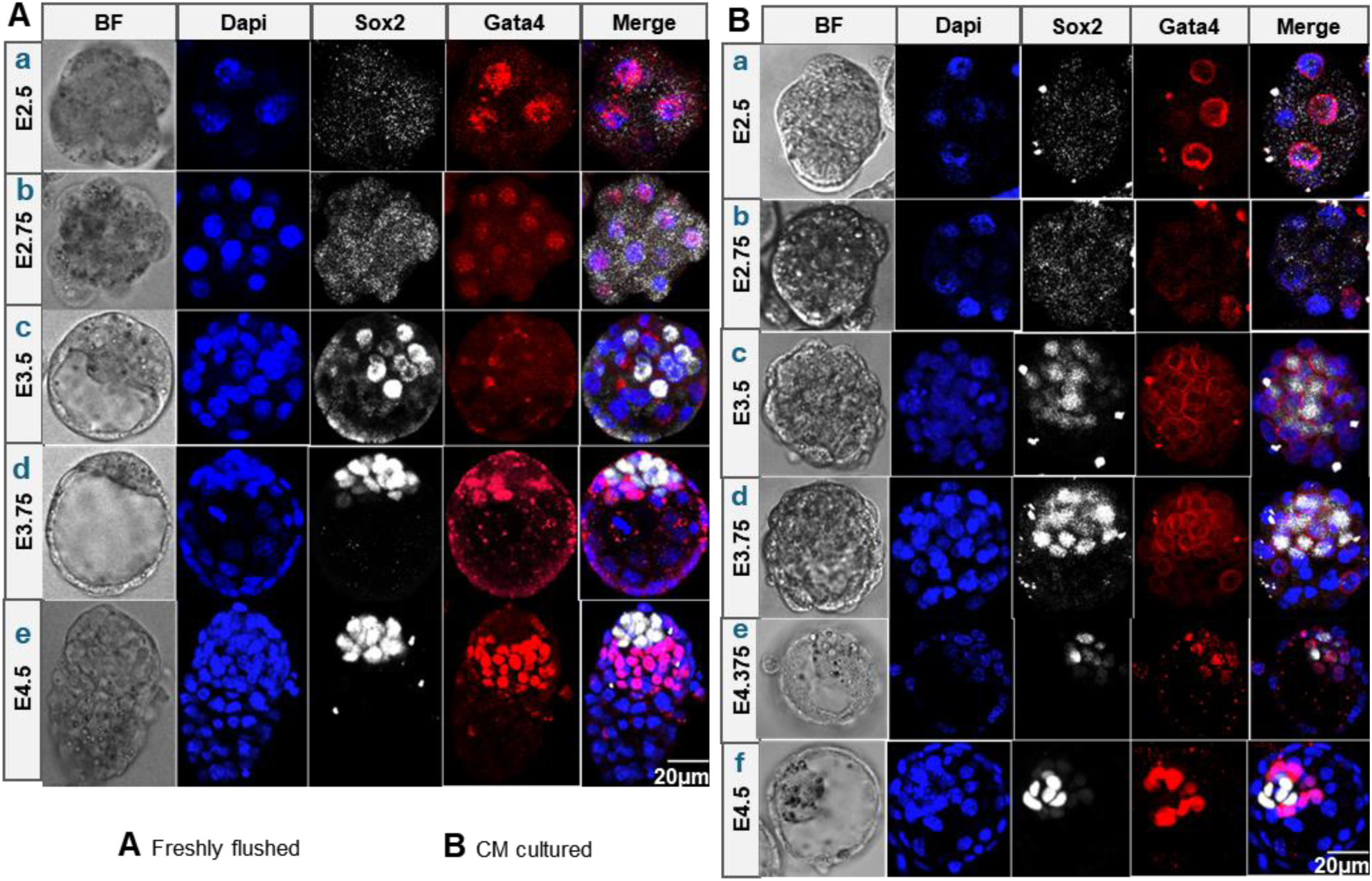
Spatiotemporal distribution of Sox2 and Gata4 expression in embryonic cells *in vivo* and *in vitro* during pre- and peri-implantation stages. **(Aa-Ae)** Sox2 and Gata4 expression *in vivo* from E2.5 to E 4.5. **(Ba-Bf)** Sox2 and Gata4 expression *in vitro* over E2.5 to E 4.5. In (A,B), DAPI-stained nuclei are shown in blue, Sox2 in white and Gata4 in red. “Freshly flushed” refers to embryos *in vivo*, and “CM cultured” refers to embryos *in vitro*. The scale bar in both (A,B): 20μm.

Moreover, at E4.5, *in-vivo* Sox2-positive cells appeared to form multiple layers, and *in-vivo* Gata4-positive cells were widely distributed, including some lining Mu-TE cells (Fig 8Ae). By contrast, at E4.5, *in-vitro* Sox2- and Gata4-positive cells tended to cluster together within restricted spatial regions within the embryos, with Sox2 remaining a limited monolayer (Fig 8Bf). Additionally, a notable difference in embryo morphology was observed between *in-vivo* and *in-vitro* groups. At E3.75, the *in-vivo* embryos appeared larger with a slightly elongated embryonic-abembryonic axis (showing a length-to-width ratio of 1.19 ± 0.08) compared to *in vitro* (having a length-to-width ratio of 1.06 ± 0.05). By E4.5, *in-vivo* embryos had significantly elongated, while *in-vitro* embryos did not exhibit the same degree of reshaping or elongation as their *in-vivo* counterparts (Fig 8Ae,Bf), in line with the results displayed earlier in this study.

### Quantitative evaluation of the dynamics of cell lineage expansion

Based on the above observed spatiotemporal differences in *in-vivo* and *in-vitro* cell lineages, I examined the quantitative trends of cell lineage emergence and expansion using the same samples as above. I focused on the dynamic changes in cell numbers of differentiated cell types, classified based on the presence of Sox2 and Gata4 markers and cell spatial location (Fig 9A). Specifically, when no cells were located inside the embryo, embryonic cells were categorised and counted only based on Sox2 and Gata4 staining. When cells were present inside the embryo but cavitation had not yet occurred, cells were classified as either internal or external. Inside cells expressing Sox2 were identified as Epi cells while those expressing Gata4 indicated PrE cells. After cavitation, external cells (TE) were categorised into Mu-TE and P-TE. The sample sizes are shown in (Fig 9B).

**Fig 9.**
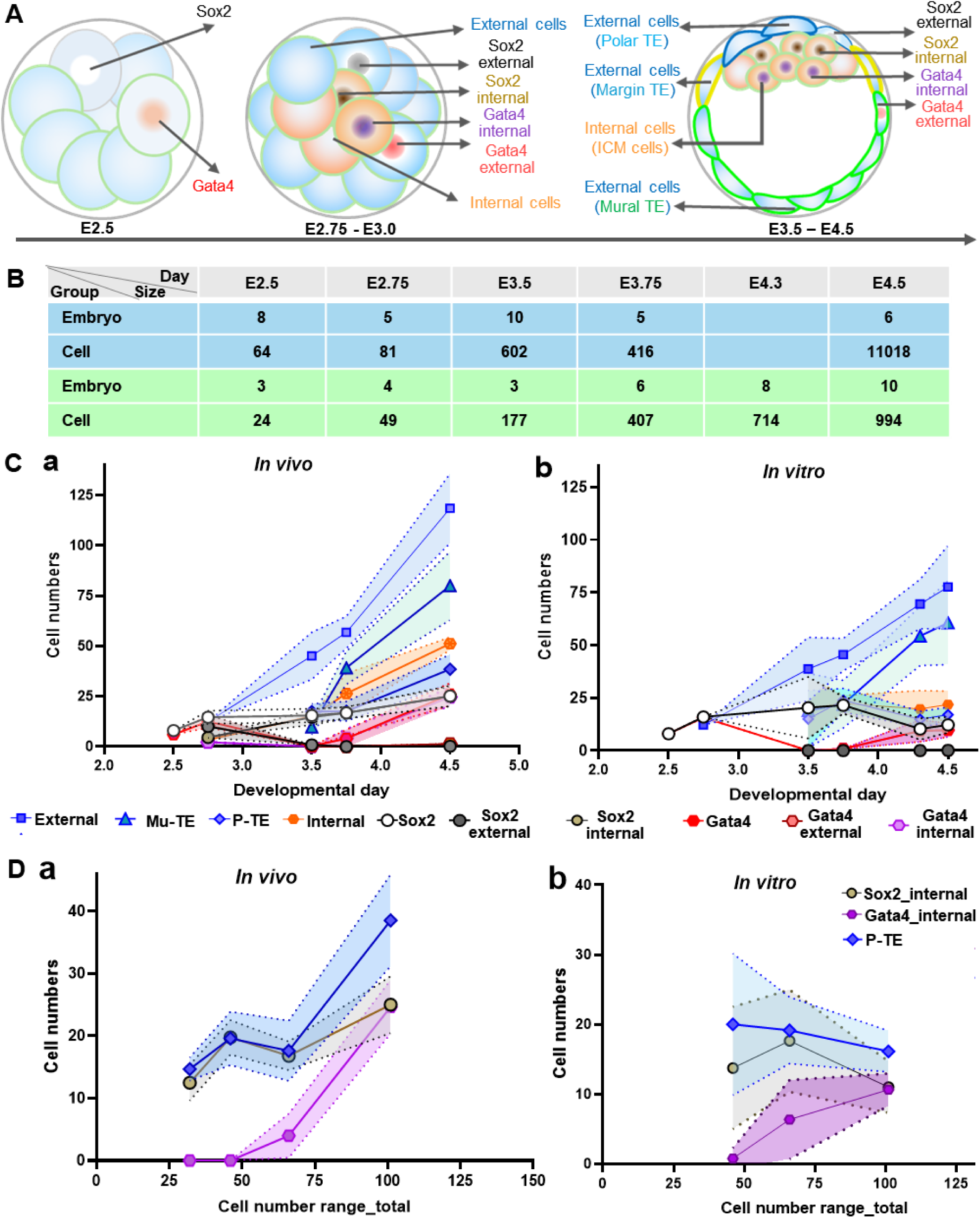
Expansion dynamics of different cell lineages/populations *in vivo* and *in vitro*. **(A)** Categorisation of different cell populations over E2.5, E2.75-E3.0, and E3.5-E4.5, respectively; Sox2-positive cells (white within nuclei), Gata4-positive cell (red within nuclei), external cells (blue), internal cell (orange), external cells with Sox2 staining (black), internal with Sox2 (brown/yellow), external with Gata4 (pink), internal with Gata4 (purple), Mu-TE (green) and P-TE (blue). **(B)** Sample sizes of embryos and cells, with blue for the *in-vivo* group and green for the *in-vitro* group, “Day” and “Size” for developmental day and sample size. *In-vivo* sample size at E4.3 (E4.275) is blank due to restricted nighttime access to the animal house. **(C,D)** Expansion dynamic curves of categorised cell populations over time; (C) for all cell population expansion over developmental days and (D) concentrating on P-TE, Gata4-ICM and Sox2-ICM lineages over cell-numbered stages. Cell populations are colour-coded corresponding to the marked colours in (A). Both *in vivo* and *in vitro* embryos were from three litters.

The results demonstrated that both *in-vivo* and *in-vitro* embryos showed a consistent increase in external cell numbers, with the *in-vivo* group tending to show a higher rate of increase, especially from E3.5 (Fig 9Cb). These trends were also observed in the Mu-TE cells of both groups and the *in-vivo* P-TE cells (Fig 9Ca,Cb). However, *in-vitro* P-TE cell numbers followed a different pattern, increasing between E3.5 and E3.75 and then dropping between E4.25 and E4.5 (Fig 9Ca,Cb). The *in-vivo* internal/ICM cells kept increasing between E2.75 and E4.5, while *in-vitro* group tended to increase more slowly between E2.75 and E3.75, then dropped at ∼E4.5 (Fig 9Ca,Cb). Similar trends were observed for total and internal Sox2-positive and Gata4-positive cell counts in both *in-vivo* and *in-vitro* groups, with *in-vitro* group showing lower rates of increase, especially between E3.75 to E4.5 (Fig 9Ca,Cb). P-TE and Sox2-positive cells, and Mu-TE and Gata4-positive cells appeared to show similar trends, respectively, in both *in vivo* and *in vitro* over time (Fig 9C,D). Employing cell numbers as an analytical benchmark of developmental timing, the results showed relatively similar trends between internal Sox2-positive cells and P-TE cells in both *in-vivo* and *in-vitro* groups (Fig 9D).

Taken together, these data show that compared to *in-vivo* embryos, *in-vitro* ones tend to follow distinct spatial and temporal patterns of Sox2- and Gata4-positive cell emergence and expansion, especially at the later stage before implantation. These findings suggest that prolonged *in-vitro* conditions alter the regulation of cell fate specification, potentially affecting the coordination of lineage segregation and organisation during pre- and peri-implantation development.

### Quantitative assessment of Epi and PrE cell lineage spatial distribution

As shown above, Gata4 cells emerged in ICM as early as E3.75 at the 67-cell stage, promoting an investigation into the specified spatial distribution of Sox2- and Gata4-positive cells before the full sorting of ICM cells. I observed that ICM surface shape varied -concave, convex, or relatively flat forms facing the cavity, while TE layer exhibited more stable morphology across embryos. To ensure consistent cell position analysis, I chose the P-TE surface centre as a reference point. An IMARIS 3D coordinate was then established with its origin aligned to P-TE apex. The positive Z-axis was oriented toward the cavity centre, while the XY plane intersected the apex of the P-TE population and remained perpendicular to the Z-axis (Fig 10B). The sample sizes of analysed embryos and cells are summarised in (Fig 10A). The result showed that, in 3D view, Gata4-positive cells tended to be already located at the top of the ICM surface before E4.5 (as early as E3.75), while Sox2-positive ICM cells were allocated under Gata4-positive cells, both *in vivo* and *in vitro* (Fig 10B,C). These findings suggest spatiotemporal differences in Sox2- and Gata4-positive cell distribution between *in-vivo* and *in-vitro* embryos, indicating that *in-vitro* condition impacts the spatial organisation of cell lineages.

**Fig 10.**
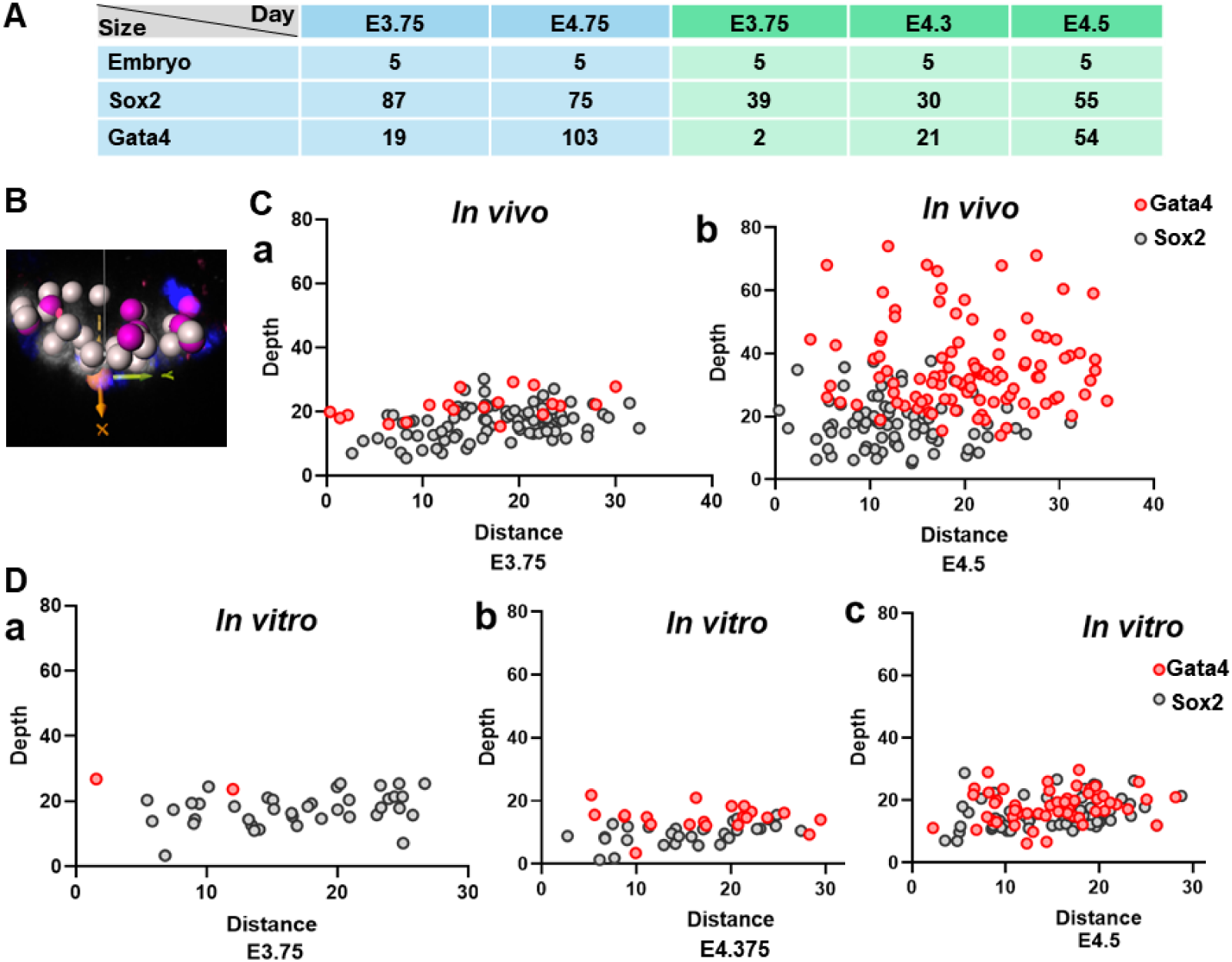
Spatial features of Gata4 and Sox2 cells *in vivo* and *in vitro* during pre- and per-implantation. **(A)** Sample sizes of *in-vivo* (blue) and *in-vitro* (green) embryos and Sox2- and Gata4-ICM cells. **(B)** Distance measurement between cells and the XY plane. The XY plane is set at the level of the P-TE apex and is oriented horizontally, perpendicular to the embryonic-abembryonic axis. The Z-axis runs parallel to the embryonic-abembryonic direction. The origin of the 3D coordinate system is at the P-TE apex, and "depth" refers to the distance of the cells from the XY plane along the Z-axis. **(Ca,Cb)** Depth of Gata4- (red) and Sox2-positive (grey) cells *in vivo* at E3.75 and E4.5, respectively. **(Dc-e)** As (C) but for E3.75, E4.375 and E4.5 *in vitro*.

## DISCUSSION

The current study compared developmental patterns between *in-vivo* and *in-vitro* conditions, particularly focusing on the spatiotemporal dynamics of morphological events and cell lineage specification. The results revealed that *in-vitro* conditions cumulatively impacted mouse embryo spatiotemporal development at multiple levels, from anatomical structures to molecule scales, including Gata4 and Sox2 expression patterns.

### Spatiotemporal Trends of Embryo Growth *In Vivo* and *In Vitro*

Cell number increases were delayed in *in-vitro* H2B-GFP mouse embryos compared to their *in-vivo* counterparts, particularly after E3.5, suggesting that the *in-vitro* micro-environment hinders embryo development. Minor differences and major similarities between *in-vivo* and *in-vitro* embryos during E1.5-E3.5 may reflect the inherent plasticity of embryos at these stages or relative similarities between the oviduct environment *in vivo* and culture medium *in vitro*. This is supported by previous studies (Hogan and Tilly, 1978; Spindle, 1978; Yao *et al*., 2019), which underscored the resilience of early-stage embryos to environmental variations. However, post E3.25/E3.5 stages, *in-vitro* conditions had a pronounced impact on embryo development in this study, potentially due to sustained application of atmospheric oxygen, laser exposure and a lack of change in culture medium, leading to the accumulation of harmful effects on *in-vitro* embryos. This receives support from previous findings showing the impacts of increased embryo susceptibility, stage-specified embryonic metabolic modes, suboptimal culture conditions, and laser exposure on *in-vitro* embryo development after E3.5 (Sakkas and Trounson, 1990; Trimarchi *et al*., 2000; Ménézo *et al*., 2013; D’Souza *et al*., 2016; Khodavirdilou *et al*., 2021). This study also showed a significant reduction in *in-vitro* embryo size, especially post-cavitation, highlighting the notable impact of *in-vitro* conditions on embryo growth and cavity expansion. This notion is supported by previous research reporting the compromised TE cell number and function in *in-vitro* human embryos, which parallels early mouse embryogenesis (Giritharan *et al*., 2012; Płusa and Piliszek, 2020).

### Spatiotemporal Patterns of Embryo Morphological Events *In Vivo* and *In Vitro*

The present work displayed that *in-vitro* compaction predominantly occurred at the 8-cell stage, consistent with previous studies (Calarco and Brown, 1969; Leivo *et al*., 1980), with occasional instances at the 4-cell stage. These two compaction patterns occurred on similar embryonic days, suggesting that while cell number generally correlates with compaction time, it does not solely dictate compaction occurrence. My findings also showed that the *in-vitro* compaction took around six hours to complete from initial cell attachment, followed by 1.5 to 2.5 hours post-completion. This indicates that compaction event is a phased process, aligning with previous studies on micro-scale cell polarisation changes during compaction (Zhu *et al*., 2017; Zhu *et al*., 2020).

My findings revealed that *in-vitro* embryos initiated cavitation at the 31- to 35-cell stages from a specific spatial site in the TE cell layer, indicating early TE cell differentiation before detection by transcriptional markers. This aligns with previously observed differences in nuclear structures between P-TE and Mu-TE cells (Ahmed *et al*., 2010). Cavitation expansion in the current study followed a non-linear pattern with rounds of volume reduction and re-expansion linked with Mu-TE cell divisions, suggesting that these divisions may trigger cavity volume adjustments. These cavity adaptive responses could stem from division-induced changes in cell-to-cell contact and cell junctions (Madan *et al*., 2007; Moriwaki *et al*., 2007).

My results showed that PrE migration *in vitro* began as early as E4.25, with initial migrating cells featuring protrusion changes, spatially arranged between certain PrE to Mu-TE cells. These findings suggest that PrE cells may interact with certain Mu-TE cells via protrusions to facilitate migration. My findings also revealed that *in-vitro* embryo hatching typically occurred shortly after cell migration initiation, mainly from Mu-TE near P-TE, a site linked to a high birth rate in ART (An *et al*., 2021). My work extends this by elucidating hatching dynamics at different TE regions. For instance, when hatching breaker cells were situated in P-TE, embryos tended to remain confined to the ZP, indicating distinctly specialised roles of P-TE, Mu-TE and their subpopulations during peri-implantation, supporting previous work on mouse embryo implantation mechanisms (Suzuki *et al*., 2022).

Compared to the discussed *in-vitro* morphological event, *in-vivo* embryos initiated compaction before E2.5, at least four hours earlier than *in vitro*. *In-vivo* cavitation began before the 31-cell stage, at least around three hours earlier than *in vitro*. In this study, *in-vivo* embryos initiated PrE cell migration as early as before E3.75, extending previous work (Christodoulou *et al*., 2019) reporting PrE migration completed between E4.0 to E4.75, providing the full view of *in-vivo* cell migration timing. In contrast, *in-vitro* embryos were still in an early stage of PrE migration as discussed above.

These results enhance our understanding of spatiotemporal dynamics of morphological events *in vitro*, while timing differences in these events between *in vitro* and *in vivo* highlight the significant impact of *in-vitro* conditions on embryo development.

### Spatiotemporal Cell Lineage Specification and Expansion *In Vivo* and *In Vitro*

My findings revealed different expansion trends of all cell lineages between *in vivo* and *in vitro*. Gata4 cells emerged as early as the 67-cell stage in *in-vivo* embryos at E3.75, with no expression in embryos with fewer than ∼67 cells from the same mouse litters, supporting previous findings on sequential cell lineage expansion (Płusa *et al*., 2008). My results suggest the 67-cell stage *in vivo* as a critical threshold for Gata4-marked cell differentiation. *In vitro*, only occasional embryos showed few Gata4 cells at the 67-cell stage around E3.75, while culturing embryos to ∼E4.5, most embryos exhibited extended Gata4 expression, even with fewer than ∼67 cells. These disparities between *in vitro* and *in vivo* highlight that cell number and developmental durations may collectively influence the timing of cell lineage-specified protein expression, consistent with hierarchical gene expression profiles (Wang *et al*., 2004; Zeng *et al*., 2004). The current study also showed that Gata4 and Sox2 cells were positioned at the ICM surface and depth from E3.75, supporting the previous findings (Gerbe *et al*., 2008) reporting that pre-PrE emerged at late E3.5. This indicates that the temporal Gata4-ICM emergence coordinates with their spatial specification as early as E3.75, expanding our understanding of the temporal Epi/PrE sorting.

My results showed that Sox2- and Gata4-cells exhibited a growth trend relatively similar to either P-TE or Mu-TE cells. These results tentatively suggest direct or indirect developmental trajectories between Epi/PrE and Mu-TE/P-TE cells. This work provides a pilot view of potential cell lineage interactions, but further investigation of cell lineage tracking and functional studies is needed to fully understand the detailed associations between various cell lineages.

### Conclusions

In summary, this study comprehensively explored spatiotemporal dynamics of embryo growth, morphological events, and cell lineage specification during pre- and peri-implantation stages, highlighting both commonalities and distinctions between *in-vivo* and *in-vitro* conditions. These insights further our understanding of influences of *in-vitro* conditions on embryogenesis, potentially causing long-term developmental issues and health problems for offspring despite *in-vitro* embryos being capable of progressing to full term. These findings also offer valuable guidance for evaluating and improving clinical practices and outcomes in ART and other fields.

### Study Limitations

In the present work, sample size at certain embryo developmental stages was limited by restricted access to the specific mouse line. Expanding the sample size in future studies could further strengthen the current conclusions. Furthermore, embryonic and cellular measurement relied on H2B-GFP marker. Future work could refine insights via the development and integration of advanced molecular-based spatiotemporal-tracing approaches with visualising cell membrane, other subcellular structures or activities while minimising technical-associated impacts on embryo development.

## MATERIALS and METHODS

### Resources and Materials

Resources and materials used in this study at the University of Manchester (UoM) are summarised in Supplementary Tables S1-S4.

### Ethical Approval and Regulations for Animal Care

Ethical approval for the animals used for this work was granted by the UoM Animal Welfare and Ethical Review Committee. The animals were bred under project license P08B76E2B, protocol 4. Animal husbandry and handling adhered to the regulations of the United Kingdom Home Office Animals (Scientific Procedure) Act 1986. All animals were humanely euthanised following Schedule 1 of the UK Animals (Scientific Procedure) Act 1986. Furthermore, all procedures followed institutional guidelines for the care and use of laboratory animals as approved by the Institutional Animal Care and Use Commitment of the UoM.

### Experimental Design

#### Workflow and parameters for investigation

This study was designed and illustrated in (Fig 11). The independent variable (IV) was the duration of *in-vitro* embryo culture and scanning, majorly categorised into two levels: freshly flushed group (*in-vivo* group), and long-term *in-vitro* group. The dependent variable (DV) contained multiple parameters associated with spatiotemporal patterning of embryogenesis over pre- and peri-implantation stages (Fig 11). Multiple parameters were considered to measure and analyse according to the experimental design, as shown in (Fig 11).

**Fig 11.**
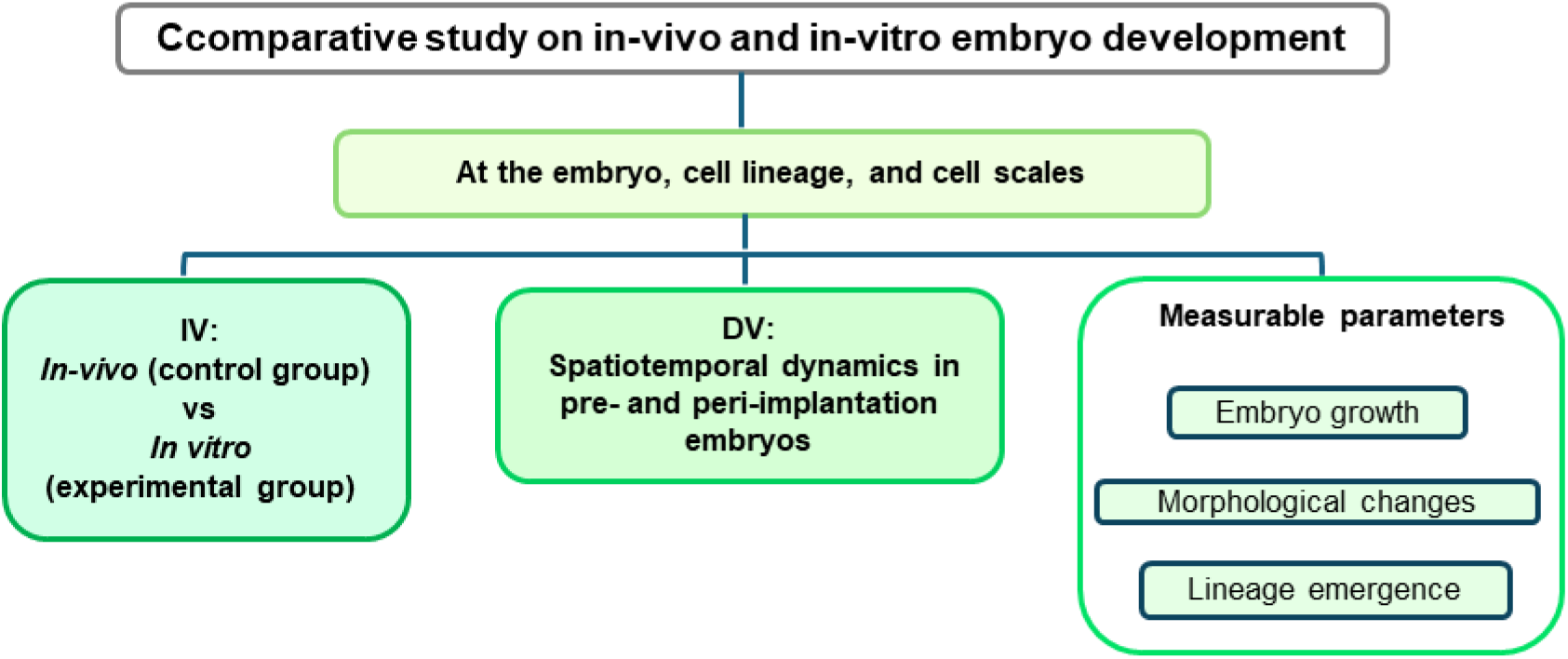
Experimental design. The top box shows the main aim of this study. The top box in light green displays the major scales investigated in the study. The left and middle dark green boxes show IVs (independent variables) and DVs (dependent variables). The right light green boxes depict main parameters measured in the study to evaluate spatiotemporal dynamic of early embryos. Green colour corresponds to the fluorescence mouse line used in the experiments.

#### Inclusive/exclusive criteria

Based on the typical development of mouse embryos and in line with my study aim, the inclusive/exclusive criteria of embryos were as follows, unless stated otherwise: (1) embryos survived and developed to the pre- or peri-implantation stages; (2) embryos displayed the stage-characterised morphology landmarks such as compaction and cavitation in the time course of interest; (3) late blastocysts exhibited proper proportions of lineages to be examined, and that was assessed through assessing both live-cell and fixed-cell images; and (4) transgenic embryos showed the visible and trackable intensity of a targeted vector.

#### Sample size

I calculated the adequate sample size based on a large effect size (Cohen’s d ≈ 1.90), supported by my initial data measurements of *in-vivo* and *in-vitro* embryonic cell number when the group difference was first detected. With 80% statistical power and a significance level of 0.05, approximately 10 embryos per group were initially targeted, which would be sufficient to detect meaningful differences. However, the sample size was adjusted to three or four embryos per group when occasionally encountering logistical constraints, such as limited embryo availability and time restrictions. Despite occasional reductions to smaller sample sizes due to resource constraints, the within-embryo repeated-measures design of my study (especially for *in-vitro* groups) enabled the detection of meaningful trends between *in-vivo* and *in-vitro* groups by taking measurements from the same embryos at multiple time points.

### Laboratory Work Procedures

#### Mice strains and maintenance

To facilitate real-time tracking of cells during embryo development, the *CAG::H2B-EGFP* transgenic mouse line, developed by Hadjantonakis Laboratory (Hadjantonakis *et al*., 2004), was employed. In this transgenic mouse line, embryonic cells express human H2B histone tagged with an enhanced green fluorescent protein (EGFP) within nuclei, allowing for chromatin visualisation. The mice used were bred and housed in the animal facility at the Building of Stopford (BSF), UoM.

The mice were maintained on a 12h:12h light-dark cycle under specific pathogen-free conditions. All female mice used for this research had a weight range from 25 g to 32 g and a roughly average age of six months. The male mice aged between six to 12 months and weighed between 32 g and 38 g. The weight of males was greater than that of females as males were reused to minimise consumption of maintained transgenic mouse strain in addition to sex dimorphism.

#### Mice mating and embryo generation

All female and male mice mated naturally, *CAG::H2B-GFPxCAG::H2B-GFP* (H2B-GFP), to produce embryos used for tracking all embryonic cells. To identify embryo development and obtain embryos on time, mating plugs were regularly checked between 9 am to 11 am every day during the experiment unless noted otherwise. The day following the visible mating plug was marked as embryonic day 0.5 (E0.5).

#### Uterus and oviduct dissection

On the targeted embryonic days, female mice were euthanised via cervical dislocation. Death was confirmed by observing the cessation of blood circulation. Afterwards, the reproductive system was dissected. The dissected uterus and oviduct were placed in a 35mmx10mm plastic dish and transported to the lab. The fat and ovary linked to the uterus and oviduct were removed in phosphate buffer solution (PBS) under binocular microscopy. Subsequently, the uterus and oviduct were moved into a home-made manipulation media (M2) dish ready for the next procedure (Grabarek and Płusa, 2012). It is noteworthy that M2 is a variation from M16 (Whittingham, 1971). The resources used here are presented in (Table S1).

#### Embryo collection and culture

Embryo flushing, collection, and transfer were conducted under a Leica stereoscope, following a protocol by Grabarek and Płusa (2012). Embryos before E3.0 were flushed from the oviducts into pre-warmed M2 medium using a glass pipette (Grabarek and Płusa, 2012). They were then transferred to pre-prepared KSOM medium (manually made in house) in a glass bottom dish, covered with paraffin oil, and incubated for 30 minutes to acquire a pH of 7.2 to 7.4 (Grabarek and Płusa, 2012). Afterwards, live embryos with a strong fluorescence signal, confirmed via the Nikon A1 confocal microscope, were selected and cultured in the KSOM dish within a controlled environmental chamber on Nikon A1 for video setup. Embryos older than E3.0 were flushed from the uterus using a 1-mL syringe. Certain embryos required the ZP removal and washing. Following ZP removal and embryo washing, the embryos with or without ZP underwent sequential procedures, including incubation and fluorescence signal checking, or fixation. Table S1 details reagents and equipment used in this procedure.

#### ZP removal

Pre-warmed AcidTyrode solution (37°C) was used to remove ZP from embryos in preparation for the fixation procedure. Embryos were gently transferred using a glass pipette into drops of pre-warmed AcidTyrode. This transfer process was repeated between sequential drops three times. Visual confirmation of ZP removal was quickly done using a Leica DM750M microscope. Following the ZP removal, embryos were promptly placed back into the M2 medium and washed several times to prepare them for subsequent procedures. For a summary of the resources used, please refer to (Table S1).

#### Fixation and immunostaining

Embryos designated for fixation were transferred to a freshly prepared solution containing 4% paraformaldehyde, 0.1% Tween20, 0.01% Triton-X, and 1X PBS. After 15 minutes, they were moved to PBS and covered with mineral oil, then stored in a refrigerator at 4°C overnight or for an extended duration, depending on the experiment schedule and equipment availability. Subsequent immunostaining was performed as previously described (Płusa *et al*., 2008). The embryos underwent two 5-minute washes in PBX, followed by permeabilisation with a medium containing 0.65% Triton-X 100 in PBS for 20 minutes at room temperature. After four times of washes, they were incubated in a blocking buffer (BB) for one hour at 4°C (see Table S2 for BB composition). Following the BB step, embryos were incubated with primary antibodies overnight at 4°C, then washed four times in PBX and blocked again with 10% donkey serum for one hour. Afterwards, they were incubated in the secondary antibody solution for 60 minutes at 4°C. Finally, embryos underwent four additional washes in PBX before nuclear staining with Hoechst in preparation for imaging. Detailed antibody information is listed in (Table S2).

#### Imaging of live and fixed embryos

Details of the equipment and software used for long videos are summarised in (Table S3). The CO_2_ incubator, which was interconnected with a Nikon A1 microscope, was checked to ensure stable conditions, including a constant temperature of 37°C and a 5% CO_2_/air mixture, and controlled humidity. The Nikon A1 was warmed up for 30 minutes before the video recording. The glass bottom dish containing the embryos was positioned on the stage of the environmental chamber within the Nikon A1. Imaging settings were selected using NIH imaging software, which were adapted from a published protocol (Płusa *et al*., 2008) (Table S4). The imaging room was kept dark during imaging. After an initial setup period of 2-3 hours, the focus of live-embryo imaging was checked, and subsequent monitoring was conducted every 12 hours to assess ongoing recording and embryo growth, based on inclusive/exclusive criteria described above. Long videos captured embryo development for approximately 69 ± 3 hours (from E1.5 to E4.5/E4.75).

Similar to the real-time movie establishment, a three-dimensional imaging acquisition of fixed embryos was carried out on A1, with certain different aspects of settings from the setup for live embryo imaging, as summarised in (Table S4).

### Image Analysis and Data Collection

#### General image analysis

Acquired ND2 files obtained from the imaging system (detailed above) were converted to IMS files via IMARIS software (Oxford company, version 9.3.1 and 9.8) for cell tracking. This work was done in the Bioimaging Facility at the UoM.

#### Counting embryonic cells

To determine temporal changes in embryonic cell numbers, I used the IMARIS spot tool in the 3D viewer to quantify nuclei/cells marked by fluorescence signals within live embryos. Additionally, cell numbers in immunostained embryos were quantified by counting cells stained for lineage markers and Dapi (Hoechst).

#### Estimating embryo size

Cell borders were identified before embryo compaction using the IMARIS 2D viewer combined with a default measurement algorithm. To further refine cell border identification at the top and bottom sections of Z-stacks, multiple IMARIS 3D visualisation tools, including Slicers and Section Viewer, were employed. On each targeted developmental day, the semi-diameter (d) of round cells, the semi-major (a), semi-intermediate (b), and semi-minor (c) axes of ellipsoid cells, were measured. Embryo size was approximated by summing the volumes of individual cells: using the formula (4/3)πd^3^ for round-shaped embryonic cells, and (4/3)π*abc for ellipsoid-shaped embryonic cells. Similarly, post-compaction, the geometric parameters of each embryo were measured, and the same formulas were applied to approximate embryo size based on their shapes: (4/3)πd^3^ for round embryos and (4/3)π*abc for ellipsoid embryos.

#### Monitoring embryo morphological changes

Embryos from freshly flushed groups were collected around developmental days E1.5, E2.5, E2.75, E3.5, E3.75, and E4.5. Due to limited access to the Animal House at the UoM during the night, it was not feasible to collect embryos between these time points. Upon collection, embryos were immediately scanned to observe cell morphological events occurring near the collection time point. For *in-vitro* cultured embryos, short longitudinal or cross-sectional images were extracted from long-term time-lapse videos to focus on each stage-specific morphological event.

#### Cavity size estimation

The volume of cavities was estimated based on embryo and cavity shapes. For cavities approximated as an ellipsoid, the volume was calculated using the formula (4/3)πabc, where a, b, and c represent the axes of ellipsoids as described earlier for evaluating embryo size. For Cavities with non-ellipsoid shapes, the volume was approximated using the formula V ≈ (4/3)πr^3^-(1/3)πr_12_(3r-r_1_), with r denoting the semidiameter of the embryo and r_1_ representing the height of the ICM cap.

#### Statistical Analysis and Data Visualisation

The data were recorded in Microsoft Excel (2016) and analysed as well as visualised using GraphPad Prism v9. Statistical analyses included: (1) Descriptive statistics were typically presented as mean ± standard deviation (SD), unless otherwise noted; (2) Independent samples between two groups of samples having non-normal distribution were analysed using a Mann-Whitney test model; (3) For multiple groups of independent samples with one IV, Kruskal–Wallis test with Dunn’s multiple comparisons test was used if data had non-normal distribution; (4) Linear regression was used for normally distributed data, while non-linear regression and curve fitting analyses were applied to non-normally distributed data, to capture complex relationships between IVs and DVs; (5) Normality of the data was assessed using P-P plot, or the Shapiro–Wilk test. When the sample size was below the minimum recommended for relevant inferential statistics, the tests were initially explored for completeness with caution and only reported when a substantial effect size and significant differences were observed. Where both the sample size and effect size were limited, only descriptive statistics or individual data points were presented. Embryo images were recorded or snapshotted on IMARIS. Schematic diagrams and illustrations were made using Inkscape and PowerPoint 2016.

## ACKNOWLEDGEMENTS

This work was derived from my self-funded PhD project. I would like to thank the colleagues in Division of Developmental Biology and Medicine, School of Medical Sciences at the UoM for reading and commenting on the related PhD thesis Chapter. I also deeply thank Bioimaging Core Facility at The UoM for their consistent and expert support in troubleshooting confocal microscopy issues and using IMARIS software. Additionally, I acknowledge the colleagues in School of Biology at the UoM for critically reviewing my manuscript and offering valuable insights. Finally, I extend my thank to the examiners for my PhD thesis and oral examination for their detailed and constructive feedback contributed to refining this work.

## DATA and RESOURCE AVAILABILTIY

All relevant data and details of resources can be found within the article and its supplementary information.

## REFERENCES

Ahmed, K., Dehghani, H., Rugg-Gunn, P., Fussner, E., et al. Global chromatin architecture reflects pluripotency and lineage commitment in the early mouse embryo. PloS One. 2010, 5, p.e10531.

An, L., Liu, Y., Li, M., et al. Site specificity of blastocyst hatching significantly influences pregnancy outcomes in mice. The FASEB Journal. 2021, 35, p.e21812.

Bertoldo, M.J., Locatelli Y., O’Neill C., Mermillod P. Impacts of and interactions between environmental stress and epigenetic programming during early embryo development. Reprod Fertil Dev. 2015, 27,1125–36.

Calarco, P.G. and Brown, E.H. An ultrastructural and cytological study of preimplantation development of the mouse. Journal of Experimental Zoology. 1969, 171, pp.253–283.

Christodoulou, N., Weberling, A., Strathdee, D., et al. Morphogenesis of extra-embryonic tissues directs the remodelling of the mouse embryo at implantation. Nature Communications. 2019, 10, 3557.

Coy P, Avilés M. What controls polyspermy in mammals, the oviduct or the oocyte? Biol Rev Camb Philos Soc. 2010, 85, 593–605.

D’Souza, F., Pudakalakatti, S.M., Uppangala, S., et al. Unraveling the association between genetic integrity and metabolic activity in pre-implantation stage embryos. Scientific reports. 2016, 6, p.37291.

Doherty, A.S., Mann, M.R., Tremblay, K.D., Bartolomei, M.S., Schultz, R.M. Differential effects of culture on imprinted H19 expression in the preimplantation mouse embryo. Biol Reprod. 2000, 62, 1526–35.

Ducibella, T. and Anderson, E. Cell shape and membrane changes in the eight-cell mouse embryo: prerequisites for morphogenesis of the blastocyst. Developmental Biology. 1975, 47, pp.45–58.

Enders, A.C., Given, R.L., and Schlafke, S. Differentiation and migration of endoderm in the rat and mouse at implantation. The Anatomical Record. 1978, 190, pp.65–77.

Fischer, S.C., Corujo-Simon, E., Lilao-Garzon, J., Stelzer, E.H., and Muñoz-Descalzo, S. The transition from local to global patterns governs the differentiation of mouse blastocysts. PLoS One. 2020, 15, p.e0233030.

Gad, A., Schellander, K., Hoelker, M., Tesfaye, D. Transcriptome profile of early mammalian embryos in response to culture environment. Anim Reprod Sci. 2012, 134, 76–83.

Gardner, D.K., Lane, M. Ex vivo early embryo development and effects on gene expression and imprinting. Reprod Fertil Dev. 2005, 17, 361–70.

Gerbe, F., Cox, B., Rossant, J., and Chazaud, C. Dynamic expression of Lrp2 pathway members reveals progressive epithelial differentiation of primitive endoderm in mouse blastocyst. Developmental Biology. 2008, 313, pp.594–602.

Giritharan, G., Piane, L.D., Donjacour, A., et al. In vitro culture of mouse embryos reduces differential gene expression between inner cell mass and trophectoderm. Reproductive Sciences. 2012, 19, pp.243–252.

Grabarek, J.B., and Płusa, B. Live imaging of primitive endoderm precursors in the mouse blastocyst. Progenitor Cells: Methods and Protocols. 2012, 916, pp.275–285.

Hadjantonakis, A.K. and Papaioannou, V.E. Dynamic in vivo imaging and cell tracking using a histone fluorescent protein fusion in mice. BMC Biotechnology. 2004, 4, pp.1–14.

Hogan, B., and Tilly, R. *In vitro* development of inner cell masses isolated immunosurgically from mouse blastocysts: II. Inner cell masses from 3·5-to 4·0-day p.c. blastocysts. Development. 1978, 45, pp.107-121.

Khodavirdilou, R., Pournaghi, M., Oghbaei, H., et al. Toxic effect of light on oocyte and pre-implantation embryo: a systematic review. Archives of Toxicology. 2021, 95, pp.3161–3169.

Kim, D.H., Ko, D.S., Lee, H.C., et al. Comparison of maturation, fertilization, development, and gene expression of mouse oocytes grown in vitro and in vivo. J Assist Reprod Genet. 2004, 21, 233–40.

Kojima, Y., Kaufman-Francis, K., Studdert, J.B., et al. The transcriptional and functional properties of mouse epiblast stem cells resemble the anterior primitive streak. Cell Stem Cell. 2014, 14, pp.107–120.

Leivo, I., Vaheri, A., Timpl, R., and Wartiovaara, J. Appearance and distribution of collagens and laminin in the early mouse embryo. Developmental Biology. 1980, 76, pp.100–114.

Madan, P., Rose, K., and Watson, A.J. Na/K-ATPase β1 subunit expression is required for blastocyst formation and normal assembly of trophectoderm tight junction-associated proteins. Journal of Biological Chemistry. 2007, 282, pp.12127–12134.

Mahdavinezhad, F., Kazemi, P., Fathalizadeh, P., et al. *In vitro* versus *In vivo*: Development-, Apoptosis-, and Implantation- Related Gene Expression in Mouse Blastocyst. Iran J Biotechnol. 2019, 17, e2157.

Ménézo, Y., Lichtblau, I. and Elder, K. New insights into human pre-implantation metabolism *in vivo* and *in vitro*. Journal Of Assisted Reproduction and Genetics. 2013, 30, pp.293–303.

Moriwaki, K., Tsukita, S., and Furuse, M. Tight junctions containing claudin 4 and 6 are essential for blastocyst formation in preimplantation mouse embryos. Developmental Biology. 2007, 312, pp.509–522.

Niwayama, R., Moghe, P., Liu, Y.J., et al. A tug-of-war between cell shape and polarity controls division orientation to ensure robust patterning in the mouse blastocyst. Developmental Cell. 2019, 51, pp.564–574.

Płusa, B. and Piliszek, A. Common principles of early mammalian embryo self-organisation. Development. 2020, 147, p.dev183079.

Płusa, B., Piliszek, A., Frankenberg, S., Artus, J., and Hadjantonakis, A.K. Distinct sequential cell behaviours direct primitive endoderm formation in the mouse blastocyst. Development. 2008, 135, pp.3081–3091.

Rinaudo P, Schultz RM. Effects of embryo culture on global pattern of gene expression in preimplantation mouse embryos. Reproduction. 2004, 128, 301–11.

Sakkas, D. and Trounson, A.O. Co-culture of mouse embryos with oviduct and uterine cells prepared from mice at different days of pseudopregnancy. Reproduction. 1990, 90, pp.109–118.

Spindle, A.I. Trophoblast regeneration by inner cell masses isolated from cultured mouse embryos. Journal of Experimental Zoology. 1978, 203, pp.483–489.

Suzuki, D., Okura, K., Nagakura, S., and Ogawa, H. CDX2 downregulation in mouse mural trophectoderm during peri - implantation is heteronomous, dependent on the YAP - TEAD pathway and controlled by estrogen-induced factors. Reproductive Medicine and Biology. 2022, 21, e12446.

Trimarchi, J.R., Liu, L., Porterfield, D.M., Smith, P.J., and Keefe, D.L. Oxidative phosphorylation-dependent and-independent oxygen consumption by individual preimplantation mouse embryos. Biology of Reproduction. 2000, 62, pp.1866–1874.

Wang, Q.T., Piotrowska, K., Ciemerych, M.A., et al. A genome-wide study of gene activity reveals developmental signaling pathways in the preimplantation mouse embryo. Developmental Cell. 2004, 6, pp.133–144.

Yang, H. Spatiotemporal Frameworks of Morphogenesis and Cell Lineage Specification in Pre- and Peri-Implantation Mammalian Embryogenesis: Insights and Knowledge Gaps from Mouse Embryo. Biology, 2025, 14, 1596.

Yao, C., Zhang, W., and Shuai, L. The first cell fate decision in pre-implantation mouse embryos. Cell Regeneration. 2019, 8, pp.51–57.

Zeng, F., Baldwin, D.A., and Schultz, R.M. Transcript profiling during preimplantation mouse development. Developmental Biology. 2004, 272, pp.483–496.

Zhu, M., Cornwall-Scoones, J., Wang, P., et al. Developmental clock and mechanism of de novo polarization of the mouse embryo. Science. 2020, 370, eabd2703.

Zhu, M., Leung, C.Y., Shahbazi, M.N., and Zernicka-Goetz, M. Actomyosin polarisation through PLC-PKC triggers symmetry breaking of the mouse embryo. Nature Communications. 2017, 8, p.921.

