## Supplemental Tables 1-4 for "Cross Sectional and Longitudinal Imaging Reveals Spatiotemporal Divergence in Morphogenesis and Cell Lineage Specification between *in-vivo* and *in-vitro* Mouse Embryo during Pre- and Peri-implantation" (from another version of this preprint: version 1, 2025-12-17)

**SUPPLEMENTARY**

**Table S1. Resources for handling fresh embryos.**

| **Apparatus** | **Elements** | **Source** |
|---|---|---|
| Surgical dissection kit | Scissor, forceps | Shared by Płusa lab |
| Cell culture dish 35mmx10mm | | Corning |
| Cell culture dish 65mmx15mm | | Corning |
| Glass bottom MatTek dish 35mmx14mm | | Mat Tek |
| Glass unplugged pipette | | Fisher |
| Syringe 1mL | Needle4.45x13mm | BD Plasticpak |
| Centrifuge tubes 15ml | | Corning Centristar |
| Filters 0.45µm | | Millex@HA |
| Pipette tips 1, 10, 200, 1000µL | | Shared by Płusa lab |
| **Equipment** | **Source** | **Identifier** |
| Microscope hot stage (-20 °C-120 °C) | SEMIC® | Shared by Płusa lab |
| Galaxy R CO2 incubator | Eppendorf | Shared by Płusa lab |
| Centrifuge | Eppendorf | 5415 R |
| Fridge (4°C) | Lec | Shared by Płusa lab |
| Lab Freezer(-20 °C) | Labcold | Shared by Płusa lab |
| Binocular stereoscope | Leica | S6 E |
| Binoculars | Leica | DM750 M |
| **Media** | **Source** | **Identifier** |
| Phosphate-buffered saline (PBS) | Gibco | 14040-091 |
| Manipulation media (M2) | Shared by Płusa lab | |
| Acidic Tyrode (AT) (pH 2.2-2.8) | Sigma Aldrich | T1788 |
| KSOM (manually made in house) | Provided by Płusa lab | |
| Mineral oil for live cells 1L | Gibco | M5310-1L |
| Mineral oil for fixed cells 500mL | Sigma | M8410 |

Note: This table lists the primary apparatus, equipment, and reagents used in the handling of freshly flushed embryos. It includes the names of these key resources, their components, sources or companies, and identifiers.

**Table S2. The resources used in embryo fixation and immunostaining.**

| Apparatus | Elements/Concentration | Source | Identifier |
|---|---|---|---|
| Graduated pasteur pipette 3ml | | Rapid electronics | 200006 |
| NunclomTM Surface | | Thermo Fisher | 176740 |
| 96-well U bottom plate | | Costar | 30319028 |
| Glass bottle 500ml | Shared by DDBM Group, FBMH, UOMA | | |
| **Equipment** | | | |
| Microwave | | Daewoo | Shared by DDBM Group |
| Denver instrument (0.1mg-60g) | | Cole-parmer | |
| **Reagents** | | | |
| Agar gel 500g | 1% Agar, 0.9%Nacl | Sigma Aldrich | A9915 |
| Sodium chloride 500g | N/A | Sigma Aldrich | S5886 |
| PBX | PBS, 0.1% Triton X-100 | Gibco | 10728775 |
| Paraformaldehyde | | Sigma Aldrich | 158127 |
| Triton X-100 | | Fluka | T8787 |
| Tween-20 | | Sigma | P1379 |
| Blocking buffer (BB) | PBS, 10% donkey serum | Sigma Aldrich | D9663 |
| Hoechst | Diluted in PBS, 1:1000 | Invitrogen | H3570 |
| **Antibodies** | | | |
| Anti-SOX2 monoclonal antibodies (I) | Diluted in BB, 1:200 | Santa Cruz Biotechnology | SC365823 |
| Goat anti-GATA4 monoclonal antibody (I) | Diluted in BB, 1:200 | Santa Cruz Biotechnology | SC1237 |
| Donkey anti-mouse antibody (II） | AlexaFlour647 | Invitrogen | A32787 |
| Donkey anti-goat antibody (II) | AlexaFlour546 | Invitrogen | A11056 |
| Donkey anti-goat antibody (II) | AlexaFlour546 | Invitrogen | A11056 |

Note: This table contains key resources for handling fixed embryos, including the primary (I) and secondary (II) antibodies. DDBM represents for Division of Developmental Biology & Medicine. FBMH stands for Faculty of Biology, Medicine and Health. UoM represents The University of Manchester.

**Table S3. Key apparatus, equipment and reagents for live-embryo imaging.**

| Apparatus | Source | Identifier |
|---|---|---|
| Glass bottle (for sterilised humidity water) | Shared by DDBM | |
| **Equipment** | | |
| Confocal microscope | Nikon | A1 |
| Temperature control system | Okalab | Shared by DDBM |
| $CO_2$/air circulation system | Shared by DDBM | |
| **Software** | | |
| NIH Elements AR | Nikon | 64-bit |
| **Agents and others** | | |
| Sterilised water | UoM | |

**Table S4. Settings used for live-embryo and fixed-embryo imaging.**

| Factors | Parameters |
|---|---|
| **Objective for video recording** | 20x/0.75NA |
| **Zoom** | 1.2-1.5 live imaging |
| | 1.7-2.0 fixed imaging |
| **Z sections** | 15-25 for live movie |
| | 32-35 for fixed embryos (depending on embryo size) |
| **Optical section distance** | 4µm live embryo imaging & all final scanning |
| | 1µm for all final scanning of live embryo imaging |
| | 1µm fixed embryo imaging |
| **Time interval** | 15/20 minutes live embryo imaging |
| | One time point fixed embryo imaging |
| **Channels** | Transmitted light |
| | Diode 405nm (only for fixed embryo imaging) |
| | GFP 488nm |
| | HeNe 546nm |
| | HeNe 633nm (only for fixed embryo imaging) |

Note: This table contains the objective, zoom, channels, and time used for live or fixed images.
